# The mitochondrial Ca^2+^ channel MCU is critical for tumor growth by supporting cell cycle progression and proliferation

**DOI:** 10.1101/2023.04.26.538295

**Authors:** Emily Fernández García, Usha Paudel, Michael C. Noji, Caitlyn E. Bowman, Jason R. Pitarresi, Anil K. Rustgi, Kathryn E. Wellen, Zolt Arany, Jillian S. Weissenrieder, J. Kevin Foskett

## Abstract

The mitochondrial uniporter (MCU) Ca^2+^ ion channel represents the primary means for Ca^2+^ uptake into mitochondria. Here we employed *in vitro* and *in vivo* models with MCU genetically eliminated to understand how MCU contributes to tumor formation and progression. Transformation of primary fibroblasts *in vitro* was associated with increased MCU expression, enhanced mitochondrial Ca^2+^ uptake, suppression of inactivating-phosphorylation of pyruvate dehydrogenase, a modest increase of basal mitochondrial respiration and a significant increase of acute Ca^2+^-dependent stimulation of mitochondrial respiration. Inhibition of mitochondrial Ca^2+^ uptake by genetic deletion of MCU markedly inhibited growth of HEK293T cells and of transformed fibroblasts in mouse xenograft models. Reduced tumor growth was primarily a result of substantially reduced proliferation and fewer mitotic cells *in vivo*, and slower cell proliferation *in vitro* associated with delayed progression through S-phase of the cell cycle. MCU deletion inhibited cancer stem cell-like spheroid formation and cell invasion *in vitro*, both predictors of metastatic potential. Surprisingly, mitochondrial matrix Ca^2+^ concentration, membrane potential, global dehydrogenase activity, respiration and ROS production were unchanged by genetic deletion of MCU in transformed cells. In contrast, MCU deletion elevated glycolysis and glutaminolysis, strongly sensitized cell proliferation to glucose and glutamine limitation, and altered agonist-induced cytoplasmic Ca^2+^ signals. Our results reveal a dependence of tumorigenesis on MCU, mediated by a reliance on mitochondrial Ca^2+^ uptake for cell metabolism and Ca^2+^ dynamics necessary for cell-cycle progression and cell proliferation.

## 1 Introduction

Calcium (Ca^2+^) transfer from the endoplasmic reticulum (ER) to mitochondria promotes bioenergetics and cell survival by stimulating the activities of dehydrogenases that control the flux of carbons through the TCA cycle, namely pyruvate dehydrogenase (PDH), isocitrate dehydrogenase (IDH) and α-ketoglutarate dehydrogenase (α-KGDH) (Denton and McCormack, 1980). ER-to-mitochondria Ca^2+^ transfer is mediated by Ca^2+^ release from the ER by inositol 1,4,5-trisphosphate receptors (InsP_3_R) and mitochondrial Ca^2+^ uptake via the mitochondrial Ca^2+^ uniporter (MCU) ion channel complex (Rizzuto et al., 1998). Under resting conditions with cytoplasmic free Ca^2+^ ([Ca^2+^]_cyt_) ∼100 nM, MCU-channel open probability is low due to a regulatory mechanism mediated by intermembrane space-localized dimeric MICU1/2 proteins, so-called channel gatekeeping (Mallilankaraman et al., 2012). Increments in [Ca^2+^]_cyt_ adjacent to the pore of an InsP_3_R channel reach concentrations >50 μM (Foskett, 2010), disabling MCU gatekeeping and promoting mitochondrial Ca^2+^ uptake. Notably, increased expression of InsP_3_R subtypes and MCU in certain cancers have emerged as features associated with aggressiveness and poor survival prognosis (Hall et al., 2014; Shi et al., 2015; Tang et al., 2015; Cardenas et al., 2016; Tosatto et al., 2016; Mound et al., 2017; Ren et al., 2017; Guerra et al., 2019; Li et al., 2020a; Li et al., 2020b; Liu et al., 2020; Wang et al., 2020; Miao et al., 2021).

We previously demonstrated that genetic or pharmacological inhibition of InsP_3_R or MCU in tumorigenic cancer cell lines and normal counterparts decreased cellular ATP content, increased NAD^+^/NADH ratio, reduced cell proliferation and activated mTOR-independent autophagy as a cell survival mechanism (Cardenas et al., 2010; Cardenas et al., 2016), emphasizing the importance of ER-to-mitochondria Ca^2+^ transfer to support basal metabolic requirements. Whereas autophagy was sufficient to enable normal cell survival, it was insufficient in cancer cells, which maintained uncontrolled proliferation leading to cell death by necrosis (Cardenas et al., 2016). Survival defects associated with acute reduction of Ca^2+^ signaling from ER-to-mitochondria could be rescued by media supplementation with nucleosides, pyruvate or α-ketoglutarate (α-KG), emphasizing compromised mitochondrial bioenergetics as the cause of the observed cancer cell death (Cardenas et al., 2010; Cardenas et al., 2020). A similar reliance on low-level constitutive ER-to-mitochondrial Ca^2+^ transfer was observed even in cancer cells with defective oxidative phosphorylation (OXPHOS) because of the Ca^2+^-dependence of TCA cycle dehydrogenases, particularly α-KGDH (Cardenas et al., 2020). Thus, matrix Ca^2+^ plays critical roles in mitochondrial bioenergetics by impinging upon OXPHOS, ATP synthesis and the flux of biochemical intermediates through the TCA cycle.

“Ca^2+^ addiction” may be a novel feature of cancer with promising therapeutic value. In support, genetic silencing of InsP_3_R enhanced apoptosis in clear cell renal cell carcinoma (Rezuchova et al., 2019) and colon cancer (Shibao et al., 2010). Genetic silencing of MCU in the breast cancer cell line MDA-MB-231 reduced cell motility and invasiveness *in vitro*, as well as tumor growth, lymph node infiltration and lung metastasis *in vivo* (Tosatto et al., 2016). Nevertheless, silencing of MCU in the same MDA-MB-231 line did not reduce cell proliferation, viability or clonogenic survival of cells exposed to different cell stressors (Curry et al., 2013; Hall et al., 2014). Destabilization of the interaction between MCU and MICU1 led to increased cell proliferation and tumor growth of lung cancer (Marchi et al., 2019), yet silencing of MICU1 in ovarian cancer cells enhanced sensitivity to cell-death stimuli and decreased cell migration (Chakraborty et al., 2017). Whereas genetic deletion of InsP_3_R in HEK293 and HeLa cells was found to generate a bioenergetic crisis associated with reduced cell proliferation, genetic deletion of MCU failed to phenocopy this effect (Young et al., 2022). Thus, the roles of ER-to-mitochondria Ca^2+^ transfer in cancer remain unclear.

Lack of genetic models has limited our understanding of the specific roles of MCU in cancer cell biology. Accordingly, we developed *in vitro* transformation models to investigate the role of MCU on the tumorigenic properties of transformed fibroblasts *in vitro* and tumor progression *in vivo*. Our results reveal a fundamental dependence of tumorigenesis *in vivo* and *in vitro* on mitochondrial Ca^2+^ uptake by MCU, mediated by a reliance on mitochondrial Ca^2+^ for cellular metabolism and Ca^2+^ dynamics necessary for cell-cycle progression and cell proliferation.

## 2 Materials and Methods

### 2.1 Cell Culture

HEK293T, HEK293T MCU-KO (generous gift from Vamsi Mootha, Harvard Medical School, Boston, MA), HEK293T MCU-rescue, immortalized fibroblasts, transformed fibroblasts and MCU-KO transformed fibroblasts were cultured in Dulbecco’s modified Eagle’s medium (DMEM, Mediatech, MT10013CM) supplemented with 10% fetal bovine serum (FBS, HyClone, SH30071.03) and 1% antibiotic-antimycotic (anti-anti, Invitrogen, 15240062), and incubated in a humidified incubator at 37°C with 95% air / 5% CO_2_. MCU-KO negative control (NC) transformed fibroblasts and MCU-rescue transformed fibroblasts were cultured in DMEM (10% FBS, 1% anti-anti) and 150 μg/mL Hygromycin B (Mediatech, MT30-240-CR), and incubated in a humidified incubator at 37°C with 95% air / 5% CO_2_.

### 2.1 Western Blotting

Cells in culture were washed with 1X Dulbecco’s phosphate-buffered saline (DPBS, Mediatech, MT21-031-CM), detached with 0.25% trypsin (Invitrogen, 15090046) and resuspended in DMEM (10% FBS, 1% anti-anti). The cell suspension was washed twice with 1X DPBS and lysed with RIPA buffer (50 mM Tris-HCl [pH 7.5], 150 mM NaCl, 1% NP-40, 0.25% deoxycholic acid, 1 mM EDTA) supplemented with 200 μM phenylmethylsulfonyl fluoride (PMSF) and protease inhibitor cocktail (Roche, 11697498001). For cell lysis, samples were placed in a tube rotator for 30 min at 4°C.

Lysates were centrifuged for 10 min at 1500 rpm. Protein concentration was determined using the Pierce BCA Protein Assay kit (Thermo Scientific, 23227). Samples were prepared for loading with 4X LICOR loading buffer (LICOR, 928-40004) and β-mercaptoethanol. Samples were boiled at 100°C for 5 min. Gel electrophoresis was performed in 4-12% Bis-Tris gels (NuPAGE, NP0322) and MOPS running buffer (Novex, NP0001). Transfer used nitrocellulose membranes (Sigma, RPN303D) in 20% MeOH Tris/Glycine buffer for 1 hr at 100V. Immunoblotted membranes were blocked with TBS Odyssey blocking buffer (LICOR, 927-50100) for 1 hr and then incubated in primary antibodies overnight at 4°C. Membranes were washed 3 times with 0.1% Tween-20 (Bio-Rad,1706531) TBS (TBTS-T) for 5 min and incubated with IRDye secondary antibodies for 1 hr in the dark. Blotted membranes were washed 3 times with TBS-T for 5 min and imaged using the Odyssey CLx system. Relative levels of MCU, PDH, pPDH, Tim23, and HSP60 were normalized to tubulin expression detected on the same blots. The antibodies used were: anti-Tubulin (1:5,000, Invitrogen, 322600), anti-MCU (D2z3B) (1:5,000, Cell signaling, 14997s), anti-Pyruvate Dehydrogenase E1-alpha subunit (1:5,000, Abcam, ab110334), anti-phospho-PDHE1-A type I (Ser293) (1:5,000, Millipore, ABS204), anti-Hsp60 (1:5,000, Abcam, ab46798), anti-Tim23 (1:5,000, BD Biosciences, 611223), IRDye 680RD goat anti-mouse (1:10,000, LICOR, 925-68070), IRDye 680RD goat anti-rabbit (1:10,000, LICOR, 926-32211), IRDye 800CW goat anti-mouse (1:10,000, LICOR, 926-68070), and IRDye 800CW goat anti-rabbit (1:10,000, LICOR, 925-3211).

### 2.2 Measurements of Mitochondrial Ca^2+^ Uptake and Membrane Potential in Permeabilized Cell Suspensions

Cells in culture were washed with 1X DPBS, detached and resuspended in DMEM (10% FBS, 1% anti-anti). The cell suspension (6×10^6^ cells in 10 mL) was incubated for 10 min in DMEM (10% FBS, 1% anti-anti) in a humidified incubator at 37°C with 95% air / 5% CO_2_, and then centrifuged for 3 min at 1000 rpm. The pellet was resuspended in Ca^2+^-free extracellular-like buffer (ECM: 20 mM HEPES-NaOH, 120 mM NaCl, 5 mM KCl, 1 mM KH_2_PO_4_, 0.2 mM MgCl_2_, 0.1 mM EGTA, pH 7.4) made using dH_2_O treated with BT Chelex® 100 resin (Bio-Rad, 143-2832) and incubated for 10 min in a humidified incubator at 37°C with 95% air and 5% CO_2_. The cell suspension was centrifuged for 3 min at 1000 rpm and resuspended in Ca^2+^-free intracellular-like buffer (ICM: 20 mM HEPES-NaOH, 10 mM NaCl, 120 mM KCl, 1 mM KH_2_PO_4_, 5 mM succinate, pH 7.5) made using dH_2_O treated with BT Chelex® 100 resin. Fluorescence was monitored in a fluorimeter with multiwavelength excitation and emission (Delta RAM, PTI) at a constant temperature of 37°C. Fura-FF (AAT Bioquest, 21028, Kd = 5.5 μM) excited at 340 nm and 380 nm was monitored at 535 nm emission. TMRE (Molecular Probes, T669) was excited at 560 nm and emission monitored at 595 nm. Addition of reagents during fluorometric measurements was performed according to the following timeline: T = 0, ICM-cell suspension; T = 25 sec, 1 μM Fura-FF and 10 nM TMRE; T = 50 sec, 0.004% digitonin; T = 100 sec, 2 μM thapsigargin (Sigma, T9033); T = 200 sec, 10 μM CGP37157 (Tocris, 1114); T = 400 sec, 3-5 μM CaCl_2_; T = 600 sec, 2 μM CCCP; T = 700 sec, 1 mM EGTA; and T = 750 sec, 1 mM CaCl_2_ (Sigma-Aldrich, 21115). To determine extramitochondrial Ca^2+^ concentration ([Ca^2+^]_cyt_) based on the ratiometric calibration of Fura-FF, we used the following equation:

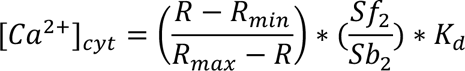

where *R* is the ratio of Fura-FF fluorescence at 340/380 nm excitation; *R_min_* is the *R* measured with [Ca^2+^] = 0; *R_max_* is the *R* measured at saturating [Ca^2+^]; *Sf*_2_ is fluorescence of Fura-FF excited at 380 nm at [Ca^2+^] = 0;. *Sb*_2_ is fluorescence of Fura-FF excited at 380 nm at saturating [Ca^2+^]; *k_d_* is the dissociation constant of Fura-FF (*k_d_* = 5.5 μM), which was experimentally validated using a set Ca^2+^ calibration buffers (World Precision Instruments, Calbuf-2). Rates of Ca^2+^ uptake were determined by fitting a single exponential from the initial peak after Ca^2+^ addition (T = 400 sec) until a steady state was reached (T = 550 sec). To determine the mitochondrial membrane potential (Δψ_m_), fluorescence of TMRE was normalized to the maximal value obtained after the addition of CCCP. Then, normalized values between T = 150 sec to T = 200 sec were averaged to obtain the reported Δψ_m_.

### 2.3 Isolation of Primary Mouse Fibroblasts

Primary mouse fibroblasts were isolated from the skin of 1-week old homozygous Mcu^fl/fl^ pups (The Jackson Laboratory, stock #029817). Pups were euthanized by decapitation. The skin was washed three times with ice-cold 1X DPBS (1% anti-anti) and sliced into 1-cm pieces using a clean razor blade. Tissue was placed in pre-warmed 500 μg/mL thermolysin solution (Sigma, T7902) prepared in HEPES buffer (Sigma, H3375) and incubated for 2 hr at 37°C with constant agitation. The tissue was transferred to a pre-warmed 0.125U/mL collagenase solution (Collagenase Type 2, Worthington Biochemical, LS004176) prepared in DMEM (10% FBS, 1% anti-anti) and incubated for 2 hr at 37°C with constant agitation. Digested tissue was forcefully ground using a 10 mL syringe plunger and centrifuged. The fibroblast-containing pellet was plated in DMEM (10% FBS, 1% anti-anti). To validate the identity of the isolated primary fibroblast population, the cells were sorted by fluorescence-activated cell sorting (FACS) for the surface marker platelet-derived growth factor receptor alpha (PDGFRA), using antibodies anti-CD140a (PDGFRA) Monoclonal Antibody (APA5) (1:100, Thermo Fisher, 14-1401-81) and goat anti-Rat IgG H&L (Alexa Fluor® 488) (1:200, Abcam, 150157), and emission filter 530 nm. HEK293T and human fibroblasts were used as negative and positive controls, respectively.

### 2.4 Generation of Lentivirus for Immortalization and Transformation of Primary Fibroblasts

Plasmids for the expression of pRRL-TERT, constitutively-active pRRL-CDK4^R24C^, dominant-negative pRRL-TP53^R248W^, and pRRL-HRas^G12V^ were generated and kindly donated by Dr. Todd W. Ridky. Lentivirus was produced in HEK293T cells. A day before transfection, HEK293T cells were seeded in 15-cm dishes, aiming for 50% confluency the next day. 24 hr after seeding, cells were co-transfected with 22.5 μg of lentiviral plasmid, 16.9 μg of the packaging plasmid psPAX2 (Addgene, 12260) and 5.6 μg of the envelope plasmid VSV-G (Addgene, 14888) using Lipofectamine 3000 (Thermo Fisher, L3000015). After 16 hr, 10 mM Na-butyrate was added into the culture. Lentivirus-containing media was collected after 48 and 72 hr of transfection and filtered through a 45 μm filter.

### 2.5 Generation of Immortalized and Transformed Primary Fibroblast Cell Lines

Primary fibroblasts isolated from 1-week old Mcu^fl/fl^ mouse pups were immortalized by sequential transduction with TERT and CDK4^R24C^ lentiviruses. Immortalized cells were kept in culture as immortalized cells or transduced with p53^R248W^ and HRas^G12V^ lentiviruses for oncogenic transformation. Genomic elimination of MCU (MCU-KO) in transformed fibroblasts was achieved by transient expression of Cre recombinase by transfection with the plasmid pLM-CMV-R-Cre (Addgene, 27546), which codes for mCherry-Cre recombinase, using Lipofectamine 3000. The cells were sorted by fluorescence-activated cell sorting (FACS) for red mCherry fluorescence. Collected cells were diluted and seeded at 1 cell/well in 96-well plates. Colonies were grown and verified by Western blot for MCU-KO. Rescue of MCU (MCU-rescue) was achieved by transfection of MCU-KO transformed fibroblasts with the plasmid pCMV3-MCU-FLAG (Sino Biological, MG5A1846-CF), which codes for MCU, using Lipofectamine 3000. Generation of MCU-KO NC transformed fibroblasts was achieved by transfection of MCU-KO transformed fibroblasts with the plasmid pCMV3-C-FLAG (Sino Biological, CV012), using Lipofectamine 3000. Transfection of MCU-KO NC and MCU-rescue was followed by Hygromycin B selection and isolation of single clones in 96-well plates. Colonies were grown and verified by Western blot for MCU expression.

### 2.6 Mouse Tumor Xenografts

All animal procedures were approved by the Institutional Animal Care and Use Committee of the University of Pennsylvania (protocol #806559). Tumor xenografts of primary mouse fibroblasts were performed in 4-8 week old male outbred athymic nude mice J:Nu (Jackson Laboratory, stock #007850). Tumor xenografts of HEK293T were performed in 4-8 week old male immunodeficient mice NOD.Cg-Prkdc^scid^/J (Jackson Laboratory, stock # 001303). Mice were housed in pathogen-free conditions in a 12 hr light/12 hr dark cycle with food and water *ad libitum*. The day of surgery, fibroblasts were washed with 1X DPBS, trypsinized and resuspended in DMEM (10% FBS, 1% anti-anti). For immortalized vs transformed xenograft experiments, cells were resuspended at a density of 10×10^6^ cells/mL. For HEK293T xenograft experiments, cells were resuspended at a density of 10×10^6^ cells/mL. For transformed vs transformed MCU-KO xenograft experiments, cells were resuspended at a density of 40×10^6^ cells/mL. Mice were anesthetized using 1.5% - 4% isoflurane in an induction chamber. Just before injection, cells were mixed with Matrigel (Fisher Scientific, 354234) in equal parts. Injection was done into the subcutaneous space of the mouse flanks. Each mouse received one injection into the subcutaneous space of one flank and another of immortalized or MCU-KO fibroblasts into the opposite flank. For HEK293T xenografts, each mouse received one injection of HEK293T WT cells into one flank and another of HEK293T MCU-KO or HEK293T MCU-rescue into the opposite flank. After injections, tumor formation and progression were monitored for 3-4 weeks and measured with a caliper. Tumor volume was determined using the following equation:

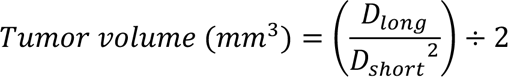

where *D_long_* is the longest diameter and *D_short_* is the shortest diameter of the tumor. At the end of the experiment, the mice were euthanized by CO_2_ exposure followed by decapitation. Tumors were excised and formalin fixed for further processing. In brief, tumors were submerged in zinc formalin solution for 24 hr at 4°C and then transferred into 70% ethanol. Tumors were embedded in paraffin (Fisher Scientific, T56-5), cut into 5-μm sections, and mounted by the Molecular Pathology and Imaging Core of the Perelman School of Medicine at the University of Pennsylvania.

### 2.7 Immunostaining and Quantification

Immunostaining was performed by the Skin Histology and Characterization Core of the Perelman School of Medicine at the University of Pennsylvania. Tumor sections were imaged at 20× using a Keyence B2-X710 microscope at 8 different regions of heterogeneous tissue excluding necrotic areas, the centers and tumor edges. Each region was imaged at 20 continuous field of views that were stitched to create a single image. Regions were imaged with three different channels: blue (DAPI), red (ki-67), and green (TUNEL) and saved as a composite. Images were imported into the software GNU Image Manipulation Program (GIMP, www.gimp.org) and the fluorescent signal of each channel was extracted as a greyscale component. Then, images were imported into the software FIJI (Schindelin et al., 2012). A signal index was determined by summing the pixel counts of the fluorescent signals as previously described (Billings et al., 2015; Asrani et al., 2019). In brief, greyscale components were transformed into binary images where the number of black pixels, representative of fluorescence, were quantified to obtain a DAPI, ki-67, and TUNEL index. Using the histogram analysis tool in FIJI, each pixel was cataloged as background or signal and then summed to obtain total number of pixels, index of background, and index of signal. As a complementary approach to quantify individual ki-67^+^ cells in tumor sections, each field of view of a region imaged with the red channel was individually processed in FIJI. First, the background was subtracted, then the image was converted into a binary component. Finally, each nucleus was represented as a single particle and counted to obtain the total number of ki-67^+^ cells.

### 2.8 Histochemistry and Histology

Tumor-section slides were processed for hematoxylin and eosin (H&E) staining by the Molecular Pathology and Imaging Core of the Perelman School of Medicine at the University of Pennsylvania. In brief, tumor sections were deparaffinized in xylene (Azer, ES609), rehydrated in ethanol (Azer, ES631) and washed with deionized water. Then, tissue was stained with hematoxylin (Leica Biosystems, 3801540) followed by eosin (Leica Biosystems, 3801600). Tissue was dehydrated with alcohol and mounted. Each slide was manually scored under the supervision of a pathologist (Dr. John T. Seykora). Necrosis analysis was done by measuring normal tissue area and necrotic area in the whole section using an eyepiece graticule with calibrated grids (25 mm). For scoring mitotic figures and giant multinucleated cells, each slide was examined at 40× at 10 different regions of heterogeneous tissue excluding necrotic areas, the centers and tumor edges. Representative images of necrotic and normal tumor tissue, mitotic figures, and giant multinucleated cells were taken at 20× using a Keyence B2-X710 microscope.

### 2.9 Sphere Formation Assay

Cells in culture were washed with 1X DPBS, trypsinized and resuspended in tumorsphere media, composed of DMEM/F12 medium (Sigma-Aldrich, D8437) supplemented with 0.4% bovine serum albumin (BSA, Life Technologies, Invitrogen, 15561020), 1% anti-anti, 20 ng/mL epidermal growth factor (Sigma-Aldrich, E5036), 10 ng/mL basic fibroblast growth factor (Sigma-Aldrich, F0291), 5 µg/mL insulin (Life Technologies, Invitrogen, A11429IJ) and 1X B27 supplement (Life Technologies, Invitrogen, 17504-044), as previously described (Johnson et al., 2013; Weissenrieder et al., 2020). Briefly, cells were plated at a density of 200 cells/well in 200 µl media in low adhesion plates, generated by applying Aggrewell solution (Stemcell Technologies, 7010) for 5 min, spinning down for 5 min at 1300 rpm and then rinsing gently with basal media (DMEM/F12, 0.4% BSA, 1% anti-anti). To reduce evaporation, outside wells were filled with sterile DPBS and not used as experimental wells. Quantification of spheroids, defined as rounded aggregates of cells with a smooth surface and poor cell-to-cell definition, was performed after 7 days of incubation.

### 2.10 Proliferation Assay

Cells in culture were washed with 1X DPBS, trypsinized and resuspended in DMEM (10% FBS, 1% anti-anti). Cells were plated at a density of 15,000 cells/well in 12-well plates at day 0. Quantification of the number of cells per well was performed for 4 days-post-seeding without continuous passage. For the proliferation assay in low nutrient conditions, cells were plated at a density of 15,000 cells/well at day 0 in high glucose (25 mM) and glutamine (4.5 mM) media. Then, at day post-seeding 1 media was changed to low glucose (4.5 mM) and glutamine (0.75 mM) media. Accordingly, quantification of the number of cells per well was performed for 4 days-post-seeding without continuous passage.

### 2.11 Transwell Invasion Assay

Cells in culture were washed with 1X DPBS, trypsinized and resuspended in DMEM (10% FBS, 1% anti-anti). Cells were plated at a density of 15,000 cells/well atop of 10 mg/mL Matrigel coated Transwells (Corning, 3464) in serum-free DMEM, while the bottom well contained 10% FBS DMEM. After 24 hr incubation, cells at the bottom of the membrane were fixed with 4% PFA (Electron Microscopy Sciences, 15713) and stained with 1 μg/μL Hoechst 33342 (Invitrogen, H3570). Quantification was done by analyzing three different fields of views of each well, imaged at 20×, and quantified automatically by the software FIJI.

### 2.12 Annexin V/DAPI FACS and Cell Viability Analysis

Cells in culture were washed with 1X DPBS, trypsinized and resuspended in DMEM (10% FBS, 1% anti-anti). Cell suspension was centrifuged at 1000 rpm for 5 min. The pellet was washed with 1X DPBS and centrifuged at 1000 rpm for 5min. The supernatant was aspirated and the cell pellet was resuspended and incubated in binding buffer with fluorochrome [488]-conjugated Annexin V (Thermo Fisher Scientific, R37174), according to manufacturer’s instructions. After 15-min incubation, 4 mM DAPI (Abcam, ab228549) was added to the suspension and cells were analyzed by FACS with emission filters 530 and 440 nm for Annexin V/DAPI detection. Data were plotted and analyzed using FCS Express software. Debris was excluded by a first gate set in forward-area and side-area (FSC-A vs SSC-A) plots. Doublets were excluded by a second level of gating in forward-area and forward-height (FSC-A vs FSC-H) plots. Cells within gates were plotted in DAPI vs Annexin V plots. Based on negative and 1 μM staurosporine-treated controls for each FACS plot, gated cells were divided into 4 quadrants: upper left (healthy cells), lower right (early apoptotic cells), upper right (late apoptotic/dead cells), and upper left (necrotic cells). Final values represent percent of gated cells in each quadrant.

### 2.13 Measurement of Oxygen Consumption Rates (OCR)

Cells in culture were washed with 1X DPBS, trypsinized and resuspended in DMEM (10% FBS, 1% anti-anti), plated at a density of 30,000 cells/well in XF96 V3 PS cell culture microplates (Agilent, 101085-004) and incubated for ∼16 hr. Media was changed to DMEM base assay medium (Sigma, D5030) supplemented with 1 mM Na-pyruvate, 2 mM glutamine, and 10 mM glucose. Baseline OCR was measured three times before the addition of 2 μM oligomycin, followed by 0.5 μM FCCP, and finally 0.5 μM rotenone/antimycin A + 1 μg/μL Hoechst (Seahorse XF Cell Mito Stress Test Kit, Agilent, 103015) according to the manufacturer’s instructions. To measure changes in OCR following acute agonist stimulation, baseline OCR determination was followed by the addition of 500 μM ATP. At the end of each assay, the center portion of each well was imaged at 20x to detect nuclei counterstained with Hoechst. Nuclei were automatically counted using FIJI and used to normalize individual OCR/well.

### 2.14 Measurement of Extracellular Acidification Rates (ECAR)

Cells in culture were washed with 1X DPBS, trypsinized and resuspended in DMEM (10% FBS, 1% anti-anti), plated at a density of 35,000 cells/well in XF96 V3 PS cell culture microplates (Agilent, 101085-004) and incubated for ∼16 hr. Media was changed to DMEM base assay medium (Sigma, D5030) supplemented with 1 mM glutamine. Baseline ECAR was measured three times before the addition of 10 μM glucose, followed by 1 μM oligomycin, and finally 50 μM 2-deoxy-glucose (2-DG) + 1μg/μL Hoechst (Seahorse XF Glycolysis Stress Test Kit, Agilent, 103020) according to the manufacturer’s instructions. At the end of each assay, the center portion of each well was imaged at 20x to detect nuclei counterstained with Hoechst. Nuclei were automatically counted using FIJI and used to normalize individual ECAR/well.

### 2.15 XTT Assay for Total Dehydrogenase Activity

Cells in culture were washed with 1X DPBS, trypsinized, resuspended in DMEM (10% FBS, 1% anti-anti) and plated at a density of 30,000 cells/well in 96-well plates. After ∼16-hr incubation, cells were incubated with XTT (Millipore Sigma, 11465015001) according to the manufacturer’s instructions. Reduction of XTT was quantified by formazan absorption at 490 nm and normalized to reference wavelength absorbance at 690 nm. XTT absorbance was detected every 30 min for 4 hrs, during which readout is linear, allowing for the best detection of differences between cell lines.

### 2.16 Measurement of Reactive Oxygen Species (ROS)

Cells in culture were washed with 1X DPBS, trypsinized, resuspended in DMEM (10% FBS, 1% anti-anti), and plated at a density of 30,000 cells/well in 96-well plates. After ∼16 hr incubation, cells were incubated with 25 μM DCFDA (Sigma-Aldrich, D6883) and 1 μg/μL Hoechst. DCFDA oxidation was detected at 535 nm and normalized to Hoechst.

### 2.17 Measurement of [Ca^2+^]_mit_

Cells in culture were washed with 1X DPBS, trypsinized, resuspended in DMEM (10% FBS, 1% anti-anti), and seeded onto coverslips. After ∼16 hr, the cells were transfected with Lipofectamine 3000 and 2.5 μg of the plasmid pCMV CEPIA2mt (Addgene, 58218). Cells were imaged 48 hr after transfection. In brief, coverslips were transferred to a perfusion chamber and perfused with extracellular-like solution (135 mM NaCl, 5.9 mM KCl, 1.2 mM MgCl_2_, 11.6 mM HEPES, 10 mM glucose, 1.5 mM CaCl_2_). Fluorescence of CEPIA2mt was monitored using a Nikon Eclipse microscope at 20x at the excitation/emission wavelengths 488 nm/500–550 nm (Suzuki et al., 2014). For each measurement, three images of the same field of view were taken 60 sec apart. Then, cells were perfused with 1 μM ionomycin (Invitrogen, 124222) followed by a 0-Ca^2+^ Tyrode’s solution (135 mM NaCl, 5.9 mM KCl, 1.2 MgCl_2_, 1 mM EGTA, 11.6 mM HEPES) to obtain maximum and minimum intensity of CEPIA2mt. Data analysis was performed using Visiview software (Visitron Systems GmbH). For each field of view, a region of interest (ROI) was delineated around the edges of each cell to obtain basal, maximum, and minimum fluorescence of CEPIA2mt. To determine the basal mitochondrial Ca^2+^ concentration ([Ca^2+^]_mit_), we used the following equation:

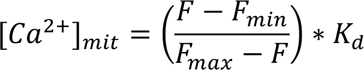

where *F* is CEPIA2mt fluorescence, *F_min_* is *F* at 0-Ca^2+^, *F_max_* is *F* at saturating Ca^2+^, and *k_d_* is the dissociation constant of CEPIA2mt (*k_d_* = 0.16 μM) (Suzuki et al., 2014).

### 2.18 Measurement of Relative Cytoplasmic [Ca^2+^]

Cells were washed with 1X DPBS, trypsinized, resuspended in DMEM (10% FBS, 1% anti-anti), and seeded onto coverslips. After 2 days, cells were incubated with 2 μM Fura-2 AM (Thermo Fisher Scientific, F1225) for 30 min at room temperature. Imaging was performed with a Nikon Eclipse Ti-U microscope using a 20x/0.75 objective. Fura-2 AM (*k_d_* = 140 nM) was excited at 340 nm and 380 nm and monitored at 535 nm emission. Before imaging, cells were perfused with 1.5 mM Ca^2+^ Tyrode’s solution (135 mM NaCl, 5.9 mM KCl, 1.2 MgCl_2_, 1.5 mM CaCl_2_, 11.6 mM HEPES) for 5 min. Baseline recording was done for 1 min while perfusing with 1.5 mM Ca^2+^ Tyrode’s solution.

Then, 2 μM ATP was added into perfusion chamber and agonist-induced responses were recorded for 300 sec. Recording was terminated after washing the cells with 0-Ca^2+^ Tyrode’s solution (135 mM NaCl, 5.9 mM KCl, 1.2 MgCl_2_, 1 mM EGTA, 11.6 mM HEPES) for 100 sec. Raw traces for each cell were obtained using Visiview. For analysis, background fluorescence recorded from a cell-free space in the field of view of each coverslip was subtracted from each image, and the background-subtracted signal was normalized to the averaged baseline value (0 – 50 sec) to obtain the ratio over time (R/R_0_). Responses were classified as oscillating, single peak, and sustained using a series of statistical parameters applied from the end of the initial peak to the end of the ATP stimulation. Those traces in which there was no agonist-induced response, the Fura-2 AM signal was significantly elevated or too low (20%), and where the responses were qualitatively identified as complex or mixed (19%), were not included in the analysis.

### 2.19 Cell Cycle Analysis

Activated lovastatin was prepared from its inactive lactone prodrug form (Millipore Sigma, PHR1285) as described (Keyomarsi et al., 1991). Cells were washed with 1X DPBS, trypsinized, resuspended in DMEM (10% FBS, 1% anti-anti), and seeded onto 6-well plates. Two days post-seeding, media of all samples, except asynchronous controls, was replaced with 15 μM lovastatin media. After 24 hr incubation, lovastatin media was replaced with DMEM (10% FBS, 1% anti-anti). At this moment, asynchronous-control and lovastatin-unreleased controls were collected and fixed. The rest of the samples were collected after 24 hr of cell cycle release from lovastatin synchronization. For sample collection and fixation, media was collected into 15 mL Eppendorf tubes, the cells were washed with 1X DPBS, trypsinized, and resuspended with the collected media. The cell suspension was centrifuged at 1000 rpm for 5 min. The supernatant was aspirated and the cell pellet was resuspended in ice-cold 1xDPBS. The cells were then centrifuged at 1000 rpm for 5 min. The supernatant was aspirated and the cells were resuspended in 100 μL of 1X DPBS. Finally, fixation buffer (80% EtOH in dH_2_O) was added in a drop-wise manner while cells were vortexed. For FACS analysis, cells were stained with 4 mM DAPI (Abcam, ab228549) and analyzed at 440 nm emission. Data were plotted and analyzed using FCS Express. Debris was excluded by a first gate set in FSC-A vs SSC-A plots. Doublets were excluded by a second level of gating in FSC-A vs FSC-H plots and verified in a DAPI-W vs DAPI-A plot. Gated cells were plotted in DAPI vs cell count histogram. Final values represent the percent of gated cells in each phase (G1, S, G2) of the cell cycle, which were determined using the Multicycle AV DNA analysis software in FCS. Unsynchronized- and lovastatin-synchronized samples were used as controls to validate the G1, S, and G2 peaks identified using Multicycle software for cell cycle analysis.

### 2.20 Measurement of Glucose and Glutamine Uptake, and Lactate and Glutamate Production

Cells in culture were washed with 1X DPBS, detached with 0.25% trypsin, and resuspended in DMEM (10% FBS, 1% anti-anti). Cells were plated at a density of 50,000 cells/well in 6-well plates in a total of 2 mL of culture media. Wells with no cells were kept as controls for normalization of metabolite concentrations in media. After 48 hrs post-seeding, media was removed, wells were washed with 1X DPBS, and fresh media was added to each well. Samples of culture media were collected 24 hrs post-media change and stored at −80°C until analysis. For normalization purposes, number of cells per well was determined manually using a hemacytometer. Quantification of glucose, lactate, glutamine, and glutamate concentrations (nM) in samples of cell culture supernatant was determined enzymatically with a bioanalyzer (YSI2950, YSI Incorporated, Yellow Springs, OH, USA). Rate of metabolite consumption (*v_c_*), rate of metabolite production (*v_p_*), cell number area under the curve (*A*), and doubling time (*d*) where calculated with the following equations:

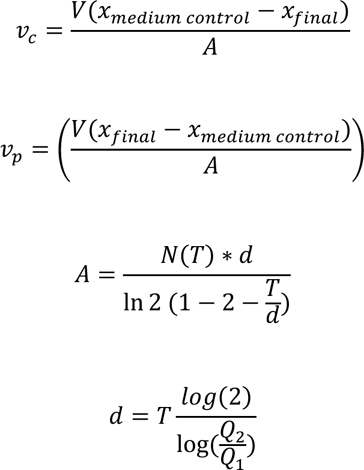

where *V* is volume of cell culture media, *x* is metabolite concentration, *A* is cell number area under the curve, *N*(*T*) is final cell count, *d* is doubling time, *T* is time of experiment, *Q*_1_ is initial cell number, and *Q*_2_ is final cell number.

### 2.21 Stable Isotope Metabolic Tracing

Cells in culture were washed with 1X DPBS, trypsinized and resuspended in DMEM (10% FBS, 1% anti-anti). Cells were plated at a density of 52,000 cells/well in 6-well plates. Tracing of labeled nutrients was performed 2 days post-seeding. For tracing of glucose carbons, no-glucose and no-glutamine media was supplemented with 5 mM of ^13^C_6_ D-Glucose (Cambridge Isotope Laboratories, Inc., CLM-1396) and 4 mM unlabeled glutamine. Tracing of glutamine was performed by supplementing no-glucose and no-glutamine media with 4 mM ^13^C_5_ L-glutamine (Cambridge Isotope Laboratories, Inc., CLM-1822-H) and 5 mM unlabeled glucose. Unlabeled controls were maintained on unlabeled low glucose (5 mM) and glutamine (4 mM) media. For metabolite extraction, after 5-hr incubation with the isotope tracers, media was removed, and cells were rinsed with 1X DPBS. Then, 500 µL of pre-chilled (on dry ice) analytical-grade 80% methanol (MeOH, Thermo Fisher, AA22909K2): 20% water (Invitrogen, 10977015) (volume/volume) was added to the well. Cell extracts were scraped and transferred to microcentrifuge tubes, and the well was rinsed with another 500 µL of 80% MeOH that was combined with the first round of extract. Samples were vortexed and incubated on dry ice for 15 min, then centrifuged at 1500 rpm at 4°C for 15 min. Supernatant containing the extracted metabolites was transferred to a new tube and stored at −80 °C until further processing. On the day of analysis, samples were dried using a vacuum concentrator (Savant SpeedVac SPD130, Thermo Fisher Scientific), and dried metabolite pellets were resuspended in 60 µL of 60:40 (v/v) acetonitrile: water, vortexed, and centrifuged at 13000 rpm at 4°C for 15 minutes. The supernatant was transferred to glass vials with polypropylene inserts for analysis by liquid chromatography-mass spectrometry (LC-MS). The instrument autosampler was maintained at 4°C, and the sample injection volume was 2.5 µL. Samples were analyzed by hydrophilic interaction chromatography coupled to a quadrupole-orbitrap mass spectrometer (Q Exactive, Thermo Fisher Scientific) via electrospray ionization. The liquid chromatography system (Vanquish Flex UHPLC with binary pump VF-P10 and split sampler VF-A10, Thermo Fisher Scientific) used a BEH amide column (ACQUITY Premier BEH Amide VanGuard FIT column, 2.1 mm x 100 mm, 1.7 µm particle size, Waters Corporation #186009508) for separation. The column was held at 35°C, and the flow rate was 300 µL/minute with a gradient of solvent A (20 mM ammonium acetate, 20 mM ammonium hydroxide in 95:5 water: acetonitrile (v/v), pH 9.5) and solvent B (acetonitrile). The gradient was 95% B to 40% B from 0 to 9 min, hold 40% B for 2 min, reverse to 95% B in 0.6 minutes, and hold at 95% B until 20 total min. Samples were directed to the mass spectrometer for 0.25 to 16 min. The mass spectrometer was operated in negative ion mode with an automatic gain control (AGC) target of 1E6, maximum inject time of 100 ms, scan range of 55-825 m/z, and 140,000 resolution. Electrospray ionization source settings included spray voltage of 3 kV and auxiliary gas heater temperature of 350°C with capillary temperature of 325°C. Raw LC-MS data were converted to mzXML file format using “MSConvert”(Chambers et al., 2012; Adusumilli and Mallick, 2017), and data were analyzed using El-MAVEN software (Melamud et al., 2010; Clasquin et al., 2012). Chemical standards for glucose, glutamine, and TCA cycle intermediates were used to validate metabolite identification. Natural ^13^C abundance correction was performed using AccuCor (Su et al., 2017).

## 3 Results

### 3.1 Transformation of Primary Mouse Fibroblasts Increases MCU Expression and Rates of Mitochondrial Ca^2+^ Uptake, Decreases pPDH Levels, and Promotes Respiration During Acute Ca^2+^-Dependent Stimulation

To investigate the roles of MCU and mitochondrial Ca^2+^ uptake in tumorigenesis, we developed *in vitro* immortalization and transformation models using a set of transgenes to immortalize and then to transform primary fibroblasts isolated from Mcu^fl/fl^ mice (Figure 1A). Isolated primary fibroblasts were transduced to overexpress TERT, the catalytic subunit of telomerase, and with constitutively-active CDK4 (CDK4^R24C^) (Figure 1A). In combination, these genes efficiently immortalize cells without conferring tumor-forming ability or genomic instability (Sasaki et al., 2009). For malignant transformation, the immortalized fibroblasts were transduced with dominant-negative p53 (p53^R248W^) and HRas (Hras^G12V^) (Figure 1A).

**Figure 1.**
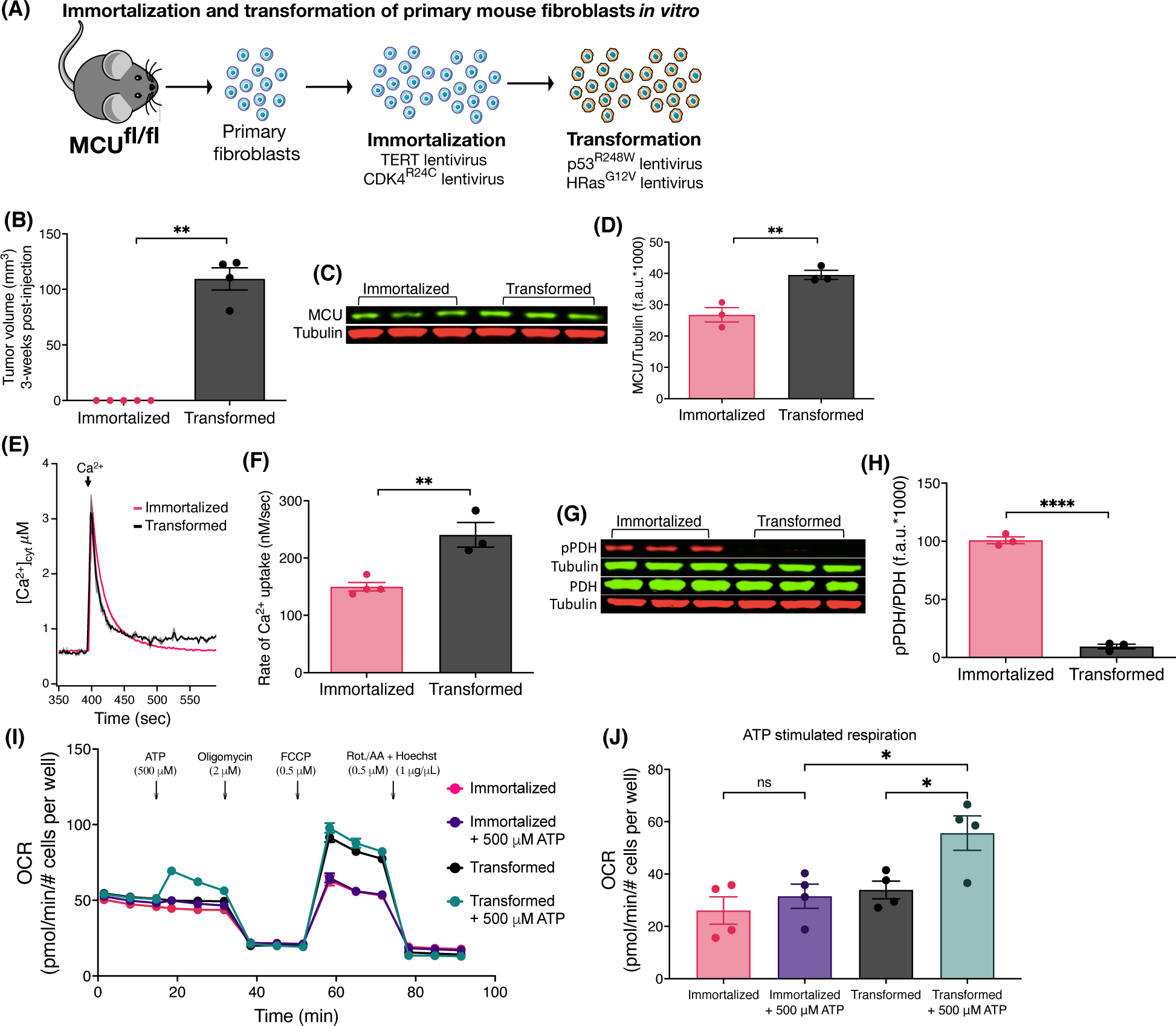
Oncogenic transformation of immortalized primary mouse fibroblasts *in vitro* induces MCU overexpression, faster rates of mitochondrial Ca^2+^ uptake, decreased pPDH levels, and promotes mitochondrial respiration during acute stimulation. (**A**) Schematic of *in vitro* immortalization and oncogenic transformation of primary mouse fibroblasts. Fibroblasts isolated from 1-week old pups of Mcu^fl/fl^ mice were transduced with the catalytic subunit of telomerase (TERT) and constitutively active CDK4 (CDK4^R24C^) in a sequential manner for immortalization. Cells were either kept in culture as immortalized cells or transduced with dominant-negative p53 (p53^R248W^) and HRas (HRas^G12V^) for oncogenic transformation. (**B**) Tumor xenograft volumes of immortalized and transformed fibroblasts 3 weeks post-injection (mean ± SEM, n = 5, **p < 0.01, Student’s t-test). (**C**) Representative immunoblots of endogenous MCU and tubulin in immortalized and transformed fibroblasts. (**D**) MCU levels normalized to tubulin expression detected on same blots. Values expressed as fluorescence units (f.a.u*1000; mean ± SEM, n = 3, **p < 0.01). (**E**) Average traces of [Ca^2+^]_cyt_ in suspensions of permeabilized transformed and immortalized fibroblasts (mean ± SEM). (**F**) Mitochondrial Ca^2+^ uptake rates of permeabilized immortalized and transformed fibroblasts (mean ± SEM, n = 3, **p < 0.01 by Student’s t-test). (**G**) Representative immunoblots of PDH, pPDH and tubulin in immortalized and transformed fibroblasts. (**H**) Relative protein levels of pPDH and PDH determined by measuring intensities of bands normalized to corresponding tubulin band intensity on same blot (mean ± SEM, n = 3, ****p < 0.0001, Student’s t-test). (**I**) Oxygen consumption rates (OCR) of transformed and immortalized fibroblasts (mean ± SEM, n = 4) in basal and stimulated conditions. (**J**) Basal and ATP-stimulated respiration in transformed and immortalized fibroblasts (mean ± SEM, n = 4, *p < 0.05, ns = non-significant, one-way ANOVA).

Subcutaneous tumors were produced by injecting immortalized and transformed fibroblasts under the skin of immunocompromised nude (J:Nu) mice. After 3 weeks post-injection, immortalized fibroblasts (n = 5) failed to form tumors whereas all injections (n = 4) of transformed fibroblasts resulted in tumors (Figure 1B). Thus, transformed fibroblasts have a malignant tumorigenic potential while immortalization serves as syngeneic control for a non-malignant phenotype.

Oncogenic mutations including Hras^G12V^ and p53^R248W^ can induce remodeling of numerous networks to support cancer progression, including mitochondrial biogenesis and turnover (Yao et al., 2019). Mitochondrial HSP60 and Tim23 expression levels were increased ∼1.5-fold in the transformed vs immortalized fibroblasts (Figure S1A), indicating that oncogenic transformation *in vitro* enhanced mitochondrial biomass (Figures S1B,C). Similarly, MCU protein expression (Figure 1C) was higher by 1.5-fold in transformed vs. immortalized fibroblasts (Figure 1D). To determine whether increased MCU expression was a transformation-associated process, we quantified MCU in three different primary fibroblast preparations (Fibroblasts WT) before any genetic manipulation (Figure S1D). WT fibroblasts expressed lower levels of MCU than transformed ones (Figure S1E). These results provide *in vitro* evidence that oncogenic transformation is associated with up-regulation of MCU. To establish the relationship between MCU expression and rates of mitochondrial Ca^2+^ uptake, cells were suspended in an intracellular-like bath solution containing the cell-impermeable Ca^2+^ indicator Fura-FF to monitor bath [Ca^2+^] ([Ca^2+^]_cyt_) and digitonin to solubilize the plasma membrane and expose mitochondria to the bath solution. Mitochondrial Ca^2+^ uptake through MCU was measured by changes in [Ca^2+^]_cyt_ after addition of a 3-5 μM bolus of Ca^2+^ to the bath solution.

Enhanced MCU expression in transformed fibroblasts was associated with significantly faster mitochondrial Ca^2+^ uptake rates compared with both untransduced WT fibroblasts (Figures S1F,G) and immortalized fibroblasts (Figures 1E,F). Enhanced mitochondrial Ca^2+^ uptake observed following transformation was not caused by altered mitochondrial membrane potential (μ4¢_m_) (not shown, but see Fig. 5C). PDH is activated by dephosphorylation by the mitochondrial matrix Ca^2+^-sensitive PDH phosphatase (PDP) (Denton, 2009). Stimulation of PDH by Ca^2+^ is known to be important for the regulation of mitochondrial metabolism and cancer progression (Denton, 2009; Pan et al., 2013; Luongo et al., 2015; Chakraborty et al., 2017; Anwar et al., 2021). Thus, we examined phosphorylation of PDH by Western blot (Figure 1G). Phospho-PDH (pPDH) was ∼4-fold lower (and almost undetectable) in the transformed cells compared with the immortalized fibroblasts (Figure 1H), suggesting that malignant transformation *in vitro* results in decreased pPDH associated with more efficient ER-to-mitochondria transfer of Ca^2+^ (Cardenas et al., 2010; Baughman et al., 2011; De Stefani et al., 2011; Hall et al., 2014; Tosatto et al., 2016; Ren et al., 2017; Yu et al., 2017; Zhao et al., 2019; Liu et al., 2020). The oxygen consumption rates (OCR) under basal and stimulated conditions were quantified to determine if oncogenic transformation resulted in alterations of mitochondrial respiration (Figure 1I). Surprisingly, cell transformation was associated with only a slight, insignificant enhancement of basal respiration (Figure S1H), although uncoupled maximal respiration was significantly increased (Figure S1I,J). More importantly, acute stimulation of mitochondrial respiration by ATP activation of metabotropic purinergic receptors promoted a rapid and significant increase of OCR in transformed, but not immortalized fibroblasts (Figure 1J). These data suggest that an increase in transformation-associated MCU expression has no effect on basal mitochondrial respiration, whereas it promotes Ca^2+^-dependent mitochondrial respiration during acute stimulation.

To further establish the relationship between MCU expression and rates of mitochondrial Ca^2+^ uptake, we utilized HEK293T cells with MCU genetically deleted (MCU-KO). In addition, these cells were used to stably express human MCU to create an isogenic HEK293T MCU-rescue line (Figure S1K). Rescue of MCU resulted in a ∼2-fold higher expression compared with WT HEK293T levels (Figure S1L). Mitochondrial Ca^2+^ uptake was absent in MCU-KO cells whereas it was restored in cells re-expressing MCU (Figure S1M). Of note, the rate of mitochondrial Ca^2+^ uptake was ∼2-fold faster in the MCU-rescue cells (Figures S1N). There were no differences between Δψ_m_ of WT, MCU-KO and MCU-rescue cells (Figure S1O), indicating that faster rates of Ca^2+^ uptake in MCU-rescue compared to WT cells were not due to differences in Δψm. These results suggest that observed higher levels of MCU expression in tumor cells are associated with enhanced mitochondrial Ca^2+^ uptake.

### 3.1 MCU is Required for Tumor Growth *in vivo* by Promoting Cell Proliferation

We developed two models to more directly explore the roles of MCU in tumorigenesis *in vivo*. In the first, we generated tumor xenografts in immunodeficient NOD SCID mice injected with WT HEK293T cells in one flank and either HEK293T MCU-KO or MCU-rescue cells in the other. When examined 4 weeks post-injection, all three cell lines had formed tumors. Compared with WT tumors, those formed by MCU-KO cells were considerably smaller (Figure S2A): WT tumors had an average volume of ∼900 mm^3^ whereas MCU-KO tumors were ∼100 mm^3^ (Figure S2B). Notably, the sizes of tumors generated by MCU-rescue cells were similar to those generated by WT cells (Figure S2B). These results suggest that MCU is dispensable for tumor formation, whereas it plays an important role in tumor growth.

In the second model, we induced tumor xenografts in immunocompromised nude (J:Nu) mice using transformed fibroblasts. We genetically eliminated MCU (MCU-KO) by expressing mCherry-Cre recombinase in the Mcu^fl/fl^-transformed fibroblasts and selecting mCherry-positive cells. Genetic deletion of MCU in clonal lines was validated by Western blot (Figure 2A) and by mitochondrial Ca^2+^ uptake assays (Figure 2B). Both cell lines formed tumors (Figure S2C), but MCU-KO tumors were significantly (> 60%) smaller (Figure 2C).

**Figure 2.**
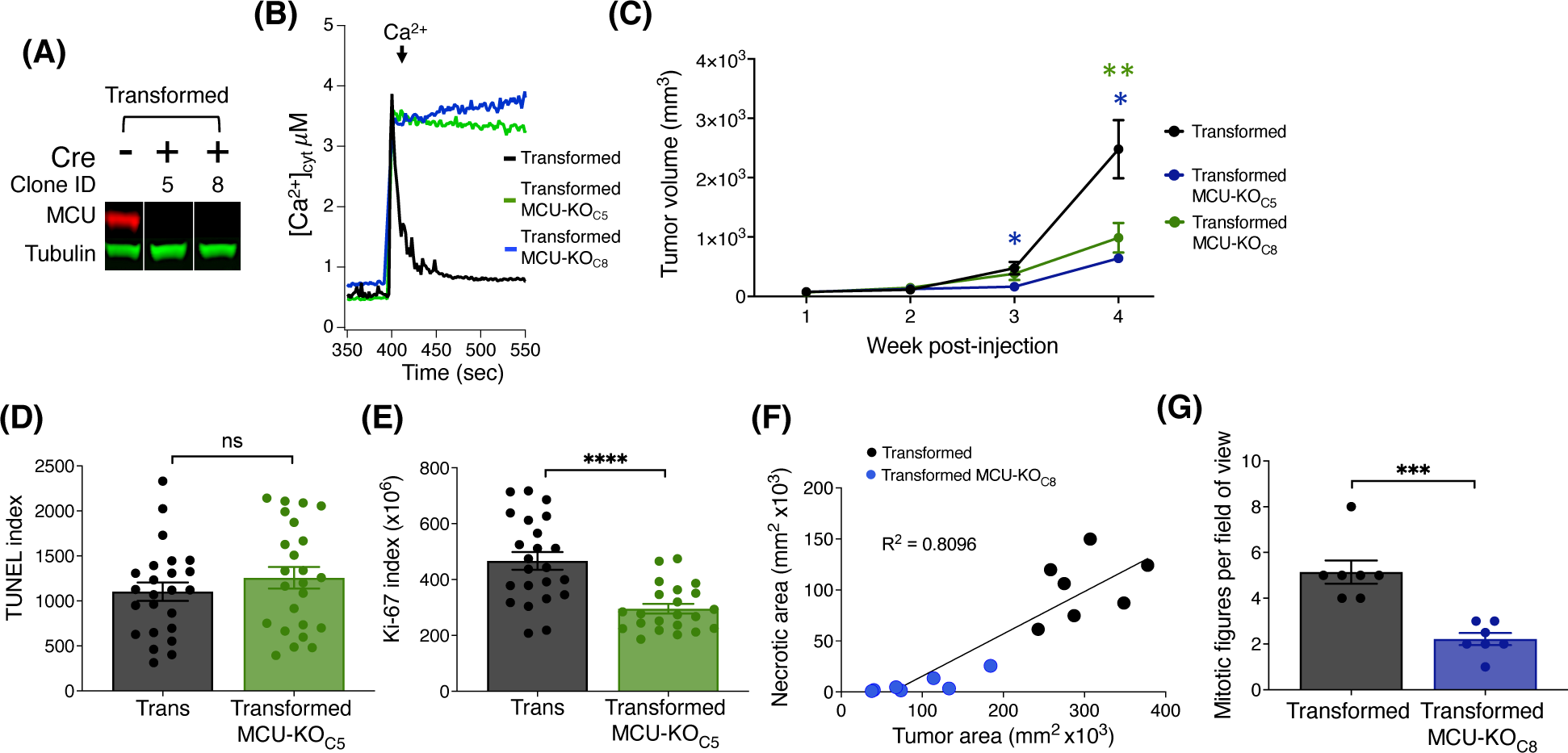
Histological analyses of tumor xenografts reveal that deletion of MCU reduces proliferation and oncogenic potential of primary mouse fibroblasts *in vivo*. (**A**) Representative immunoblots of MCU and tubulin in transformed fibroblasts. Single MCU-KO clones selected from heterogeneous population of transformed fibroblasts expressing mCherry-Cre by FACS. (**B**) Representative traces of [Ca^2+^]_cyt_ in cell suspensions of transformed MCU-KO clones to validate complete elimination of MCU-mediated mitochondrial Ca^2+^ uptake. (**C**) Volumes of tumor xenografts of transformed fibroblasts and MCU-KO fibroblasts during 4 weeks post-injection (mean ± SEM, n = 10, * p < 0.05, ** p < 0.01, two-way ANOVA). (**D**) Quantification of TUNEL index in tumor xenografts as indicator of DNA damage due to cell death (mean ± SEM, n = 3, ns = non-significant, Student’s t-test). (**E**) Quantification of ki-67 index in tumor xenografts as indicator of cell proliferation (mean ± SEM, n = 3, ****p < 0.0001, Student’s t-test). (**F**) Necrotic area relative to total tumor area. Necrotic and normal areas quantified manually. (**G**) Quantification of mitotic figures in H&E-stained xenograft sections. For each tumor, 10 different fields of view examined at high magnification (40x) (mean ± SEM, n = 7, ***p < 0.001, Student’s t-test).

To explore the mechanisms underlying reduced tumor size associated with lack of MCU, we quantified cell death and the proliferation index of fibroblast tumor xenografts. Paraffin-embedded tumor slices were stained with DAPI, immunolabeled for ki-67 as a marker of cell proliferation, and TUNEL (terminal deoxynucleotidyl transferase-mediated dUTP Nick-end labeling)-stained as a marker of apoptotic cell death (Figures S2D). Notably, cell death was not enhanced in tumors formed by MCU-KO transformed fibroblasts compared with those formed by MCU-expressing transformed fibroblasts (Figure 2D). In contrast, the proliferation index of MCU-KO tumors was markedly lower than in the tumors of transformed fibroblasts (Figure 2E). To validate this, we analyzed small tumor-tissue sections and counted individual ki-67^+^ cells (Figures S2E). The proliferation potential of transformed tumor fibroblasts was strikingly decreased by elimination of MCU (Figure S2F). These results suggest that MCU-KO does not eliminate the tumorigenic potential of transformed fibroblasts, whereas it markedly slows tumor growth primarily by strongly reducing cell proliferation with lesser enhancement of cell death.

To further understand the mechanism by which deletion of MCU impedes tumor growth, we performed histological analyses of tumors (Figure S2G), examining characteristics associated with tumor growth patterns, including amount of normal and necrotic tumor tissue (Figure S2H) and mitotic activity (Figure S2I). Necrosis in tumors often indicates aggressiveness associated with high cell density due to rapid cell division with low availability of nutrients and anoxic conditions. In agreement, the smaller tumors formed by MCU-KO transformed fibroblasts were associated with significantly reduced size of necrotic areas (Figures 2F). The number of mitotic figures was considerably lower in MCU-KO tumors (Figure 2G), in agreement with the ki-67 analysis. Some cells with morphological characteristics associated with senescence, namely markedly large cell size and/or polyploidy, were recognized as giant multinucleated cells (Figure S2J). Of interest, a substantially higher number of multinucleated giant cells was observed in MCU-KO tumors (Figure S2K).

### 3.2 Deletion of MCU Reduces Cell Proliferation of Primary Mouse Fibroblasts

The results from the *in vivo* transformed fibroblast tumor model suggests that MCU is required for cell proliferation to support tumorigenesis. We undertook a series of *in vitro* experiments to evaluate the effects of MCU depletion on cell proliferation, as well as other cellular phenotypes that drive cancer cell malignancy, including inhibition of cell death, cell division, sphere formation and matrix invasion. The transformed fibroblasts proliferated significantly faster than the immortalized fibroblasts, as expected (Figure 3A). Notably, proliferation of the transformed fibroblasts was only moderately reduced by genetic deletion of MCU in both clonal MCU-KO lines examined (Figure 3A). MCU-KO mice have no apparent physiological phenotypes until they are physically stressed (Kwong et al., 2015; Luongo et al., 2015). Therefore, we examined proliferation of transformed fibroblasts under low-nutrient conditions, a stress imposed in the tumor microenvironment. Fibroblasts were initially seeded in high glucose (25 mM) and glutamine (4.5 mM) media, and after 24 hrs the media was changed to one with low glucose (4.5 mM) and glutamine (0.75 mM) that was sufficient to allow fibroblasts to proliferate and remain viable for a period of 4 days post-seeding. Under these conditions, genetic deletion of MCU much more strongly decreased proliferation of transformed fibroblasts (Figure 3B). To confirm the requirement of MCU expression for optimal cell proliferation, we generated MCU-rescue cell lines, which were validated by Western Blot (Figures S3A,B) and by mitochondrial Ca^2+^ uptake assays (Figures S3C,D). Proliferation was enhanced in both rescue cell lines in both nutrient-rich (Figure S3E) as well as in nutrient-poor (Figure S3F) conditions, confirming that MCU expression correlates with cell-proliferative capacity and ruling out the possibility that decreased proliferation of MCU-KO cells was due to non-specific effects of Cre expression.

**Figure 3.**
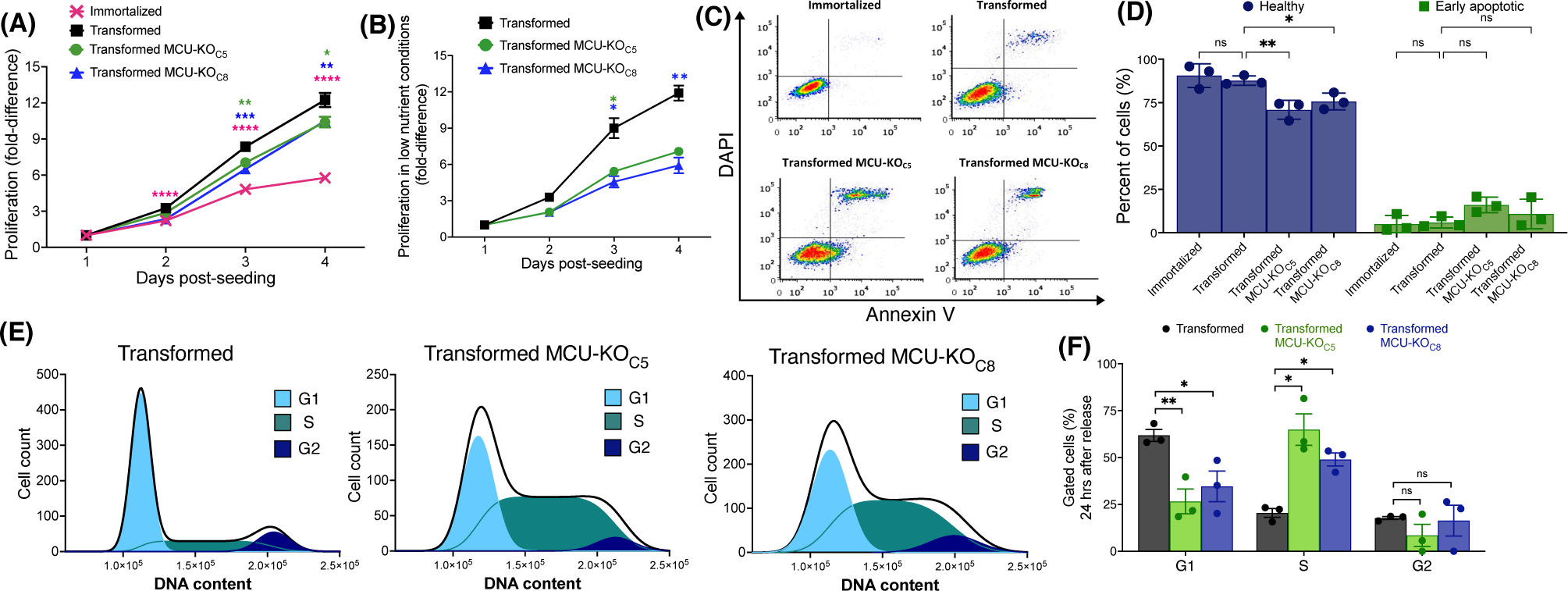
Deletion of MCU reduces proliferative potential, invasion ability, and delays cell-cycle progression of transformed mouse fibroblasts *in vitro*. (**A**) Proliferation of immortalized and transformed fibroblasts. Each data point represents 3 biological replicates in triplicate. Fold-difference represents number of cells normalized to cell count at day 1 post-seeding (mean ± SEM, n = 3, *p < 0.05, **p < 0.01, ****p < 0.0001, two-way ANOVA compared to transformed cells). (**B**) Proliferation of transformed fibroblasts in low-nutrient conditions. Each data point represents 4 biological replicates in triplicate. Fold-difference represents number of cells normalized to day 1 post-seeding. (mean ± SEM, n = 4, *p < 0.05, **p < 0.01, two-way ANOVA compared to transformed cells). (**C**) Representative FACS plots of Annexin-V/DAPI-stained cells. (**D**) Percent (%) healthy cells (lower-left quadrant, purple) and early apoptotic cells (lower-right quadrant, green) of each individual FACS plot significantly different in (C), (mean ± SEM, n = 3, *p < 0.05, **p < 0.01, ns = non-significant, two-way ANOVA compared to transformed cells). (**E**) Representative histograms of DNA-content distribution of transformed fibroblasts collected after 24 hrs of release from lovastatin synchronization. Samples were fixed, stained with DAPI, and FACS sorted for quantification. (**F**) Quantification of percent (%) of gated cells in G1, S, and G2 after 24 hrs release from lovastatin synchronization for transformed fibroblasts (black bars), MCU-KO clone 5 (green bars), and MCU-KO clone 8 (blue bars) (mean ± SEM, n = 3, *p < 0.05, **p < 0.01, ns = non-significant, two-way ANOVA compared to transformed cells).

To determine the mechanisms of reduced proliferation of transformed cells lacking MCU, we examined cell death and cell-cycle progression. Apoptotic and non-apoptotic cell death were assessed by annexin-V/DAPI staining and fluorescent-activated cell sorting (FACS) (Figure 3C). Populations of live, dead, apoptotic, and non-apoptotic cells were established based on viable and 1 μM staurosporine-treated controls (Figure S3G). There were no differences in the percent of either non-viable (early/late apoptotic and necrotic cells) or healthy cells between the immortalized and transformed populations (Figure S3H). Genetic deletion of MCU diminished the percentage of healthy cells and moderately increased the number of early-apoptotic cells (Figure 3D). Thus, the major effect of MCU-KO in transformed cells was to decrease cell proliferation. Notably, these *in vitro* results are highly consistent with the conclusions reached in the *in vivo* model.

To understand the basis for reduced proliferation observed *in vivo* and *in vitro*, we examined the effects of MCU deletion on cell-cycle progression. Fibroblasts were synchronized in the G1 phase using lovastatin (Keyomarsi et al., 1991), and then examined by DAPI-FACS 24 hr after release from synchronization. Unsynchronized and lovastatin-synchronized samples provided controls (Figure S3I). At 24 hrs after release from synchronization, a significant fraction of the transformed cells progressed through mitosis into the G2 phase of the cell cycle (Figures 3E,F). In contrast, both MCU-KO clonal cell lines contained a significantly lower percent of cells in G1 phase and a much higher percentage in S phase (Figure 3E,F). Accumulation of MCU-KO cells in the S phase suggests that mitochondrial Ca^2+^ uptake is important for cell proliferation by promoting progression through the cell cycle (Cardenas et al., 2016; Koval et al., 2019; Zhao and Pan, 2021).

### 3.3 Deletion of MCU Reduces Cancer-Associated Phenotypes of Primary Mouse Fibroblasts

The ability to form clonally-derived spheres on non-adherent substrates is a stem cell-like capacity related to metastatic tumor initiation and progression (Uchida et al., 2010). In a sphere-formation assay, the transformed fibroblasts, but not the immortalized cells, readily formed spheres (> 30 spheres per well) (Figure 4A), consistent with their differential abilities to form tumors *in vivo* (Figure 1B). In contrast, MCU-KO transformed fibroblasts formed < 5 spheres per well (Figure 4A). These results suggest that MCU plays an important role in both cell proliferation as well as the capacity to self-renew.

**Figure 4.**
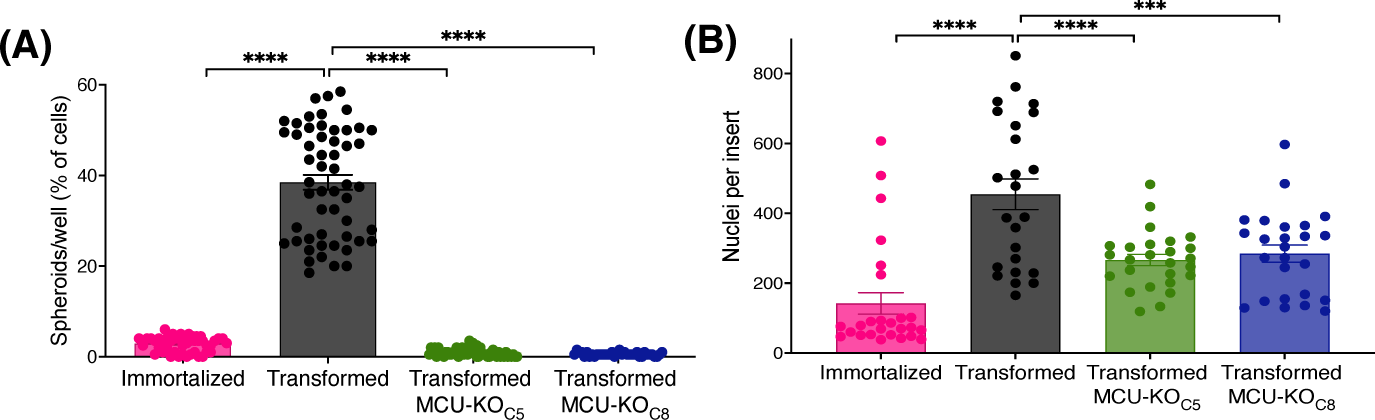
MCU deletion inhibits sphere formation capacity and reduces cell invasion *in vitro*. (**A**) Number of spheroids formed as a percent of number of cells seeded. 200 cells were plated in each well of low adhesion plates and spheroids were counted 7 days later (mean ± SEM, n = 4, ****p < 0.0001, one-way ANOVA compared to transformed cells). (**B**) Transwell invasion assay. 15,000 cells were seeded atop Matrigel-coated Transwells. After 24 hrs, membranes were fixed and stained with Hoechst 33342 to quantify number of invading cells. Three different fields of view of each well were imaged and quantified (mean ± SEM, n = 3, ****p < 0.0001, one-way ANOVA compared to transformed cells).

To explore the role of MCU in cell chemotaxis and extracellular matrix invasion, processes associated with cancer metastasis in which Ca^2+^ signaling has been proposed to play a role (Tang et al., 2015; Tosatto et al., 2016), we employed a Transwell-invasion assay. Cells were seeded atop a porous Matrigel-coated membrane and examined after 24 hr (Figure S4A). The number of transformed fibroblasts that penetrated through the coated membrane was ∼3-fold greater than the number of immortalized fibroblasts that traversed it (Figure 4B). Importantly, genetic deletion of MCU in transformed fibroblasts markedly reduced the number of invading cells by ∼50% (Figure 4B).

Together, these results validate our transformed fibroblasts as a model that recapitulates many phenotypes associated with malignancies, including enhanced proliferation, ability to form clonally-derived spheres, and cell migration and tissue invasion. Importantly, elimination of MCU-mediated mitochondrial Ca^2+^ uptake strongly suppressed these *in vitro* phenotypes, suggesting that it could play an important role in carcinogenesis, as observed in our *in vivo* models.

### 3.4 Deletion of MCU in Transformed Primary Mouse Fibroblasts Increases Glycolysis

To investigate mechanisms by which elimination of MCU-mediated mitochondrial Ca^2+^ uptake affects cancer-progression phenotypes, we examined several mitochondrial functions. First, we examined total mitochondrial dehydrogenase activity using the XTT assay. Oncogenic transformation significantly increased dehydrogenase activity, but genetic deletion of MCU was without effect (Figure 5A). Similarly, ROS production was decreased after transformation but was not changed by deletion of MCU (Figure 5B). ι¢14¢_m_ was not significantly different after transformation or in MCU-KO fibroblasts (Figure 5C). In addition, mitochondrial matrix [Ca^2+^] ([Ca^2+^]_mit_) was also not different after transformation or in MCU-KO cells (Figure 5D). Together these findings indicate that reduced malignancy of MCU-KO transformed fibroblasts is not associated with reduced mitochondrial function.

**Figure 5.**
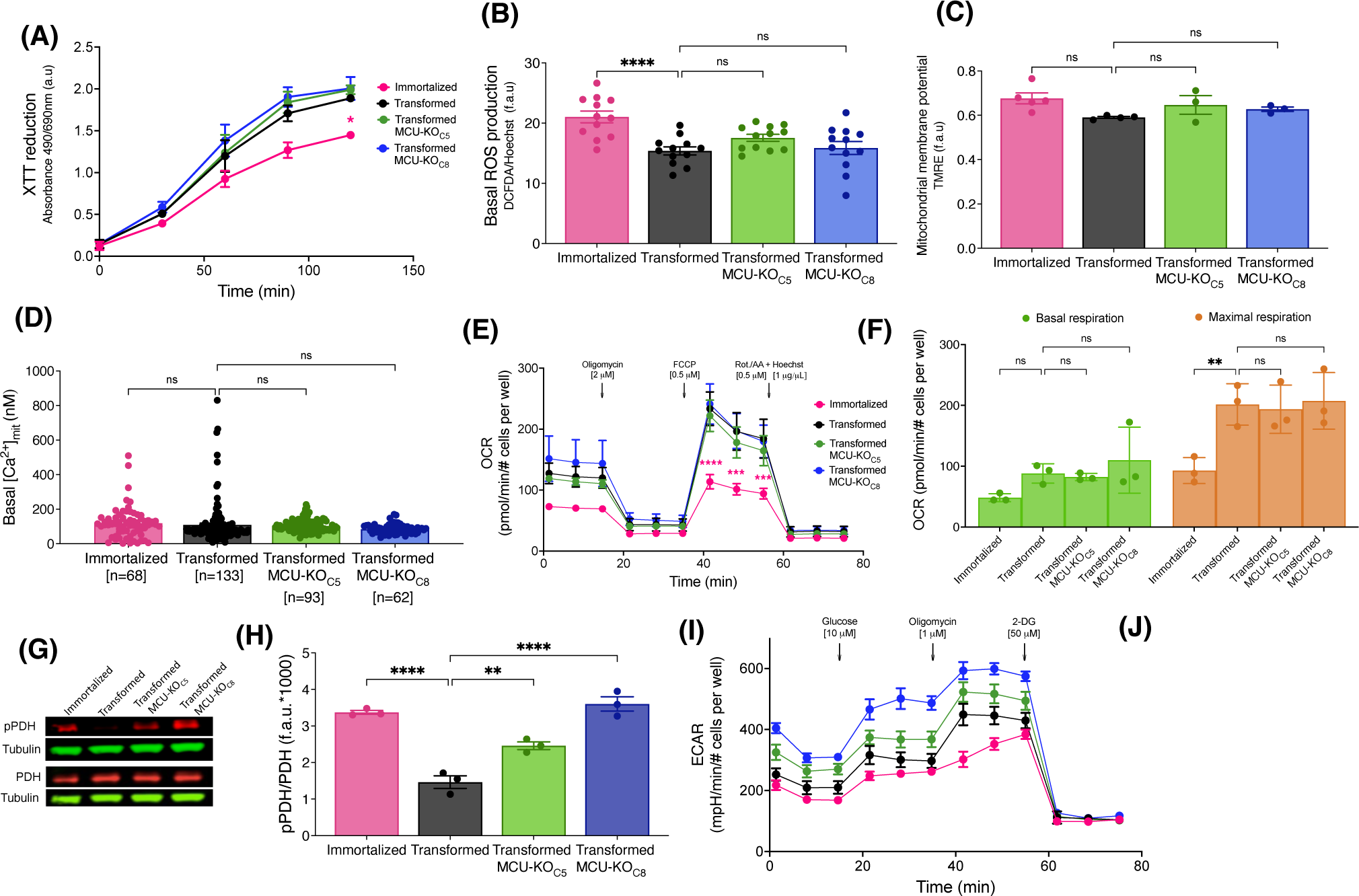
Cell physiological effects of MCU deletion in transformed primary mouse fibroblasts *in vitro*. (**A**) Metabolic activity measured by XTT reduction (mean ± SEM, n = 3, *p < 0.05, two-way ANOVA compared to that of transformed fibroblasts). (**B**) Basal reactive oxygen species (ROS) production measured over 45 min by DCFDA fluorescence normalized to Hoechst 33342 (mean ± SEM, n = 3, ****p < 0.0001, ns = non-significant, one-way). (**C**) Normalized Δψ_m_ measured by TMRE fluorescence (f.a.u) in permeabilized cell suspension (mean ± SEM, n = 3, ns = non-significant, one-way ANOVA). (**D**) Resting mitochondrial matrix [Ca^2+^] ([Ca^2+^]_mit_) measured with the indicator CEPIA2mt. Each cell was individually analyzed (n = number of cells; mean ± SEM, ns = non-significant, one-way ANOVA). (**E**) OCR of transformed and immortalized fibroblasts. Mean values compared to those of transformed fibroblasts (mean ± SEM, n = 3, ***p < 0.001, ****p < 0.0001, two-way ANOVA compared to transformed fibroblasts). (**F**) Basal (green bars) and maximal (orange bars) respiration of transformed and immortalized fibroblasts (mean ± SEM, n = 3, **p < 0.005, ns = non-significant, two-way ANOVA). (**G**) Representative immunoblots of PDH, pPDH and tubulin in transformed and immortalized fibroblasts. (**H**) Relative protein levels of pPDH and PDH determined by measuring intensities of bands normalized to corresponding tubulin band intensity on same blot (mean ± SEM, n = 3, **p < 0.01, ****p < 0.0001, one-way ANOVA). (**I**) Glycolytic extracellular acidification rates (ECAR) of immortalized and transformed fibroblasts (mean ± SEM, n = 4). (**J**) Basal glycolytic ECAR of immortalized and transformed fibroblasts. For statistical analysis, mean values were compared to that of transformed fibroblasts (mean ± SEM, n = 4, *p < 0.05, ****p < 0.0001, ns = non-significant, one-way ANOVA).

To further explore the role of MCU in cellular bioenergetics, we quantified OCRs (Figure 5E). Unexpectedly, neither basal nor maximal respiration of transformed fibroblasts was affected by genetic deletion (Figure 5F) or rescue (Figures S5A,B) of MCU. Spare respiratory capacity, proton leak, coupling efficiency, ATP production, and non-mitochondrial OCR were also not altered by transformation, MCU-KO or MCU-rescue (data not shown). Nevertheless, diminished phosphorylation of PDH associated with transformation (Figure 2G) was strongly suppressed by genetic deletion of MCU (Figures 5G,H) and enhanced by MCU-rescue (Figures S5C,D). Although genetic deletion of MCU did not affect OCR, it significantly increased glycolysis (Figures 5I,J) and glycolytic capacity (Figure S5E). The glycolytic reserve of fibroblasts was increased after transformation and remained unchanged after MCU-KO (Figure S5F). No changes were observed in non-glycolytic acidification rates after transformation or deletion of MCU (data not shown). Thus, the major effects of MCU deletion on cellular bioenergetics *in vitro* was enhanced PDH phosphorylation and upregulation of glycolysis, while mitochondrial respiration remained unaffected.

### 3.5 Deletion of MCU in Transformed Primary Mouse Fibroblasts Alters Cellular Metabolism

Altered metabolism is a fundamental characteristic of cancer cells (Hanahan and Weinberg, 2011). Cancerous cells metabolize large quantities of glucose into lactate to support rapid energy and biomass production for cellular growth and proliferation (Hatzivassiliou et al., 2005; DeBerardinis et al., 2008a; Deberardinis et al., 2008b; Lunt and Vander Heiden, 2011; Ganapathy-Kanniappan and Geschwind, 2013; Liberti and Locasale, 2016). Indeed, the observed increase in ECAR in MCU-KO cells was associated with increased glucose uptake (Figure 6A) and lactate production (Figure 6B), effects that were attenuated by MCU re-expression (Figure 6A,B). When the rate of lactate production was normalized to glucose uptake rate, there were no significant difference between cell lines (Figure 6C), suggesting that glucose is largely metabolized to lactate through glycolysis in MCU-KO cells. To determine if increased glucose consumption in MCU-KO cells alters cellular metabolism, we performed metabolic tracing with ^13^C_6_ D-Glucose (Figure 6D). The network depicted in Figure 6D represents metabolic routes, labeling pattern of ^13^C_6_ D-Glucose-derived metabolites, and fractional labeling of isotopologues of interest. Glucose entry into the TCA cycle via acetyl-CoA and PDH leads to labeling of TCA cycle intermediates with two ^13^C, represented by m+2.

**Figure 6.**
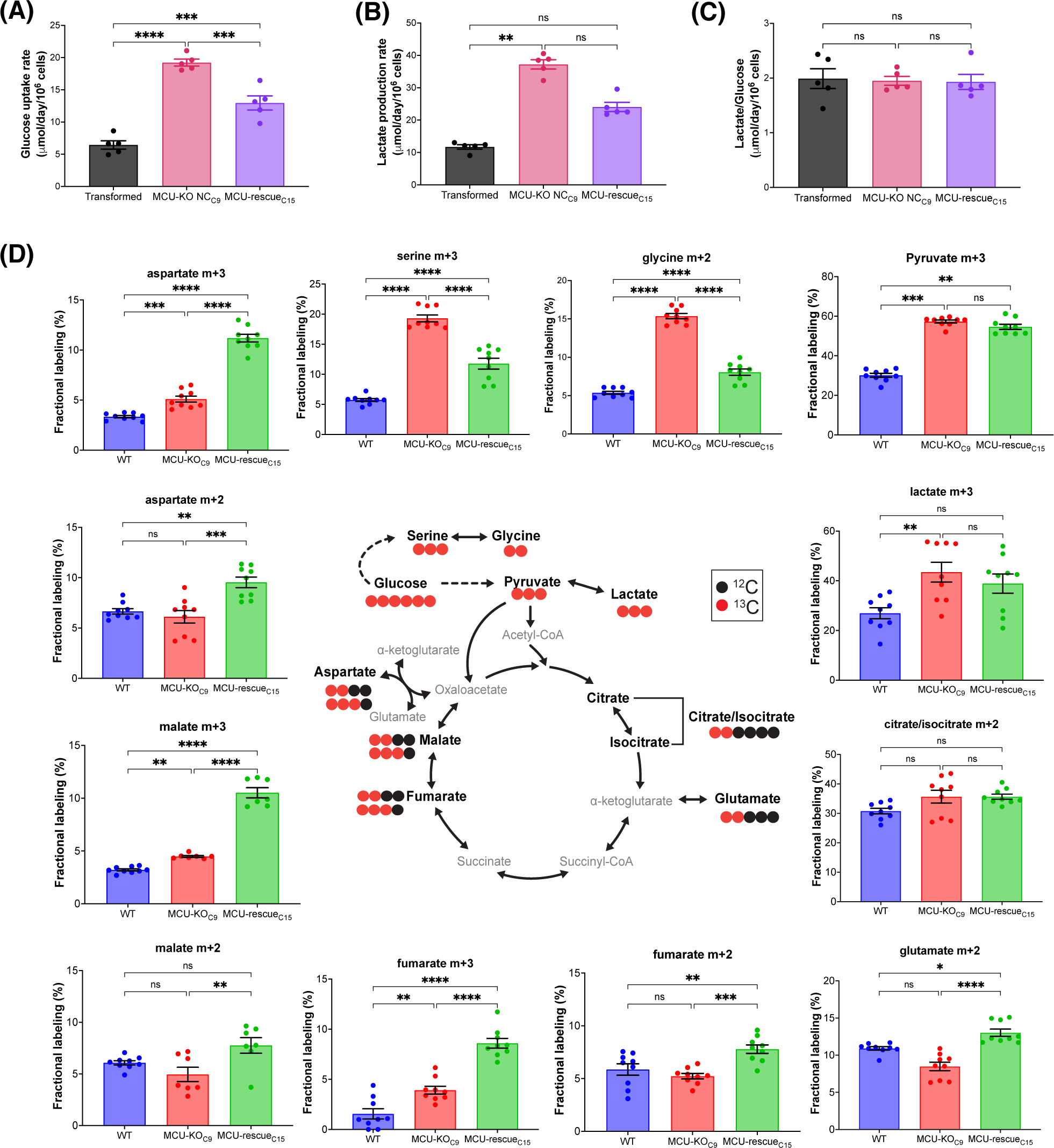
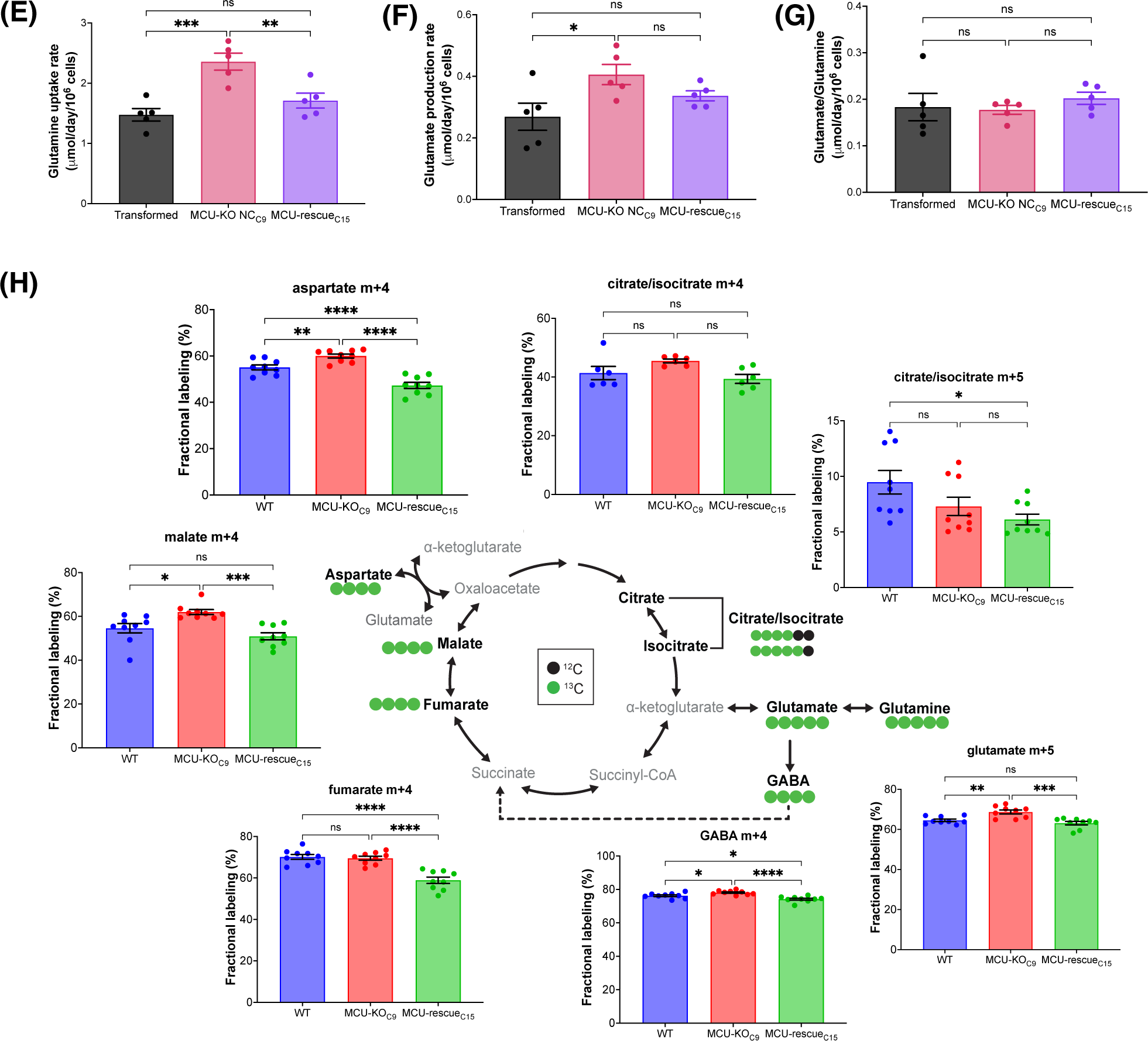
Deletion of MCU alters glucose and glutamine metabolism. (**A**) Glucose-uptake rates of transformed fibroblasts (mean ± SEM, n = 5, ***p < 0.001, ****p < 0.0001, one-way ANOVA). (**B**) Lactate-production rates of transformed fibroblasts (mean ± SEM, n = 5, **p < 0.01, ns = non-significant, one-way ANOVA). (**C**) Lactate-production rates normalized to glucose-uptake rates (mean ± SEM, n = 5, ns = non-significant, one-way ANOVA). (**D**) Schematic representation of metabolic routes, labeling pattern of ^13^C_6_ D-Glucose-derived metabolites, and fractional labeling of isotopologues of interest (mean ± SEM, n = 8, ns = non-significant, *p < 0.05, **p < 0.01, ***p < 0.001, ****p < 0.0001, one-way ANOVA). Red circles represent labeling with ^13^C and black circles represent naturally-occurring ^12^C. Isotopologues of interest are represented by mass (m) and total number of ^13^C. (**E**) Glutamine-uptake rates of transformed fibroblasts (mean ± SEM, n = 5, ns = non-significant, **p < 0.01, ***p < 0.001, one-way ANOVA). (**F**) Glutamate-production rates of transformed fibroblasts (mean ± SEM, n = 5, ns = non-significant, *p < 0.05, one-way ANOVA). (**G**) Glutamate-production rates normalized to glutamine-uptake rates (mean ± SEM, n = 5, ns = non-significant, one-way ANOVA). (**H**) Schematic representation of metabolic routes, labeling pattern of ^13^C_5_ L-Glutamine-derived metabolites, and fractional labeling of isotopologues of interest (mean ± SEM, n = 8, ns = non-significant, *p < 0.05, **p < 0.01, ***p < 0.001, ****p < 0.0001, one-way ANOVA). Green circles represent labeling with ^13^C and black circles represent naturally occurring ^12^C.

Alternatively, entry of glucose-derived ^13^C into the TCA cycle via pyruvate carboxylation by pyruvate carboxylase (PC) or malic enzyme (ME) results in the incorporation of three ^13^C in TCA-cycle intermediates, represented by m+3. Labeling of lactate in aerobic glycolysis (m+3) was significantly increased in MCU-KO as compared to transformed cells (Figure 6D). Notably, labeling of m+3 serine and m+2 glycine was also increased in transformed MCU-KO cells. This phenotype was reversed by MCU-rescue (Figure 6D), suggesting that deletion of MCU promotes the diversion of glucose-derived carbons into biosynthetic pathways, such as for purine biosynthesis (Vanhove et al., 2019). Interestingly, m+2 glutamate, fumarate, malate, and aspartate levels were not different between transformed and MCU-KO cells, although rescue of MCU enhanced their labeling beyond those observed in the wild-type transformed cells (Figure 6D). This result is consistent with the observed lack of effect of MCU deletion on basal OCR. Notably, elevated levels of m+3 fumarate, malate, and aspartate indicate that entry of pyruvate through alternative pathways, likely PC, was significantly enhanced by MCU-KO (Figure 6D). These results suggest a striking metabolic adaptation in MCU-KO cells that promotes serine biosynthesis and TCA cycle activity.

Many cancer cells shift their substrate preference to fuel flux through the TCA cycle (Reitzer et al., 1979). Glutamine provides another key carbon source for the TCA cycle through anaplerosis (DeBerardinis et al., 2007; Wise et al., 2008). We therefore evaluated the contribution of glutamine to the TCA cycle. The schematic in Figure 6H represents the labeling pattern of ^13^C_5_ L-Glutamine-derived TCA cycle intermediates, γ-aminobutyric acid (GABA), and fractional labeling of isotopologues of interest. Isotopic labeling with ^13^C is represented by green circles, meanwhile naturally-occurring ^12^C is depicted as black circles. Glutamine anaplerosis by oxidation of α-KG results in a m+4 mass increase of TCA-cycle intermediates. On the other hand, partial reverse flow of the TCA cycle for reductive carboxylation of α-KG by enzymatic activity of Ca^2+^-independent IDH2 leads to formation of citrate-isocitrate m+5. Genetic deletion of MCU was associated with an increased glutamine uptake (Figure 6E) and its conversion to glutamate (Figure 6F). With glutamate production normalized to glutamine uptake rate, there was no difference between cell lines (Figure 6G), suggesting that MCU deletion was primarily associated with increased nutrient uptake rather than involvement of alternative glutamine metabolic pathways. In agreement, ^13^C_5_-glutamine tracing revealed increased labeling of m+5 glutamate, m+4 aspartate, and m+4 malate in MCU-KO vs WT transformed cells, which was reversed by MCU re-expression (Figure 6H). We also found a significant increase in the diversion of glutamine-derived carbons into the GABA shunt in MCU-KO cells and significant decrease with MCU-rescue (Hoang et al., 2021). The GABA shunt can serve as a reservoir for TCA cycle anaplerosis by promoting entry of glutamine-derived carbons through succinate and by-passing α-KGDH (Figure 6H). Taken together, these data suggest that in the absence of MCU-mediated mitochondrial Ca^2+^ uptake, glucose metabolism via glycolysis is enhanced, and cells rely more on glutamine to maintain TCA-cycle integrity.

### 3.6 Deletion of MCU in Transformed Primary Mouse Fibroblasts Alters Agonist-Induced Cytoplasmic Ca^2+^ Signals

Because mitochondria can play an important role in buffering changes of [Ca^2+^]_cyt_, and alterations in [Ca^2+^]_cyt_ may regulate cell-biological functions of tumor cells, we also examined InsP_3_R-mediated Ca^2+^ signaling in transformed fibroblasts and two clones with MCU deleted. Exposure of cells to ATP elicited 3 types of responses: a single transient spike, oscillations, and a sustained rise (Figure 7A). In transformed fibroblasts, 60% of cells responded with a sustained elevation of [Ca^2+^]_cyt_, 35% displayed [Ca^2+^]_cyt_ oscillations, and 4% responded with a single [Ca^2+^]_cyt_ spike (Figure 7B). In contrast, a sustained elevation was rarely observed in transformed fibroblasts lacking MCU, with cells responding with either single spikes (∼41%) or oscillations (∼56%) (Figure 7B). The amplitude of the first [Ca^2+^]_cyt_ peak was elevated in the cells lacking MCU (Figure 7C). Among the oscillatory responses (Figure 7D), those in the cells with MCU deleted were of lower frequency compared with those of transformed fibroblasts (Figure 7E). Taken together, these results suggest that [Ca^2+^]_cyt_ signaling is altered in transformed cells by deletion of MCU.

**Figure 7.**
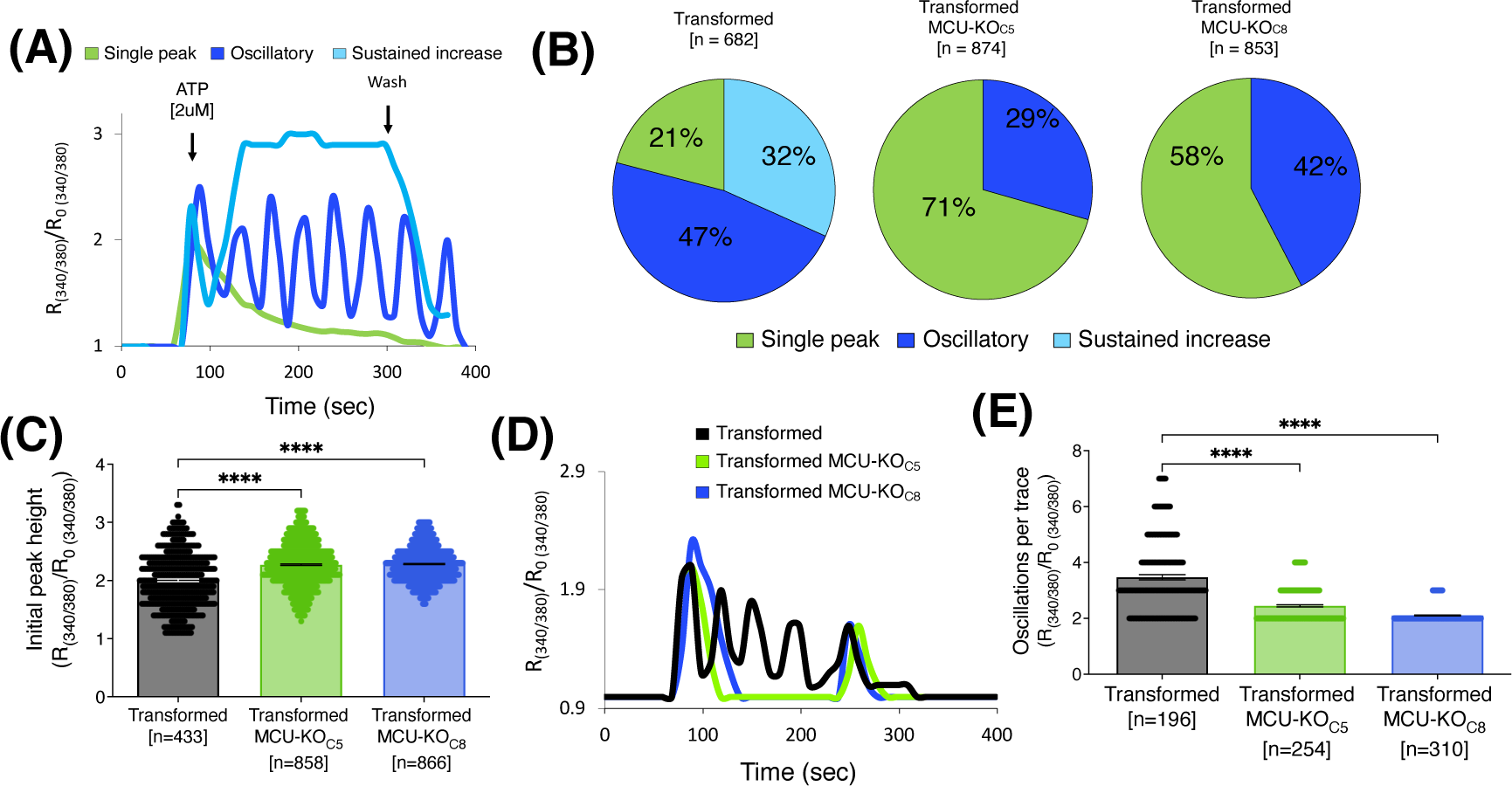
Deletion of MCU alters cytosolic Ca^2+^ transients induced by activation of metabotropic purinergic receptors by ATP. (**A**) Representative traces of relative cytoplasmic [Ca^2+^] measured with Fura-2 and represented as R/R_0_ for normalization, where R is the ratio of Fura-2 fluorescence at 340:380 nm excitation. ATP-induced responses in transformed fibroblasts classified as single spike (green trace), oscillations (blue trace), or sustained rise of [Ca^2+^]_cyt_ (light blue trace). (**B**) Percent (%) of ATP-induced responses classified as single spike (green), oscillatory (blue), and sustained rise of [Ca^2+^]_cyt_ (light blue) in transformed fibroblasts and MCU-KO fibroblasts. Each cell was individually analyzed (n = number of cells). (**C**) Height of initial peak R/R_0_ after addition of ATP (n = number of cells; mean ± SEM, ****p < 0.0001, one-way ANOVA). (**D**) Representative traces of oscillatory responses induced by ATP in transformed (black trace), MCU-KO clone 5 (green trace), and MCU-KO clone 8 (blue trace) fibroblasts. (**E**) Number of oscillations over 300 sec in responses classified as oscillatory (n = number of cells; mean ± SEM, ****p < 0.0001, one-way ANOVA).

## 4 Discussion

In the present study, we examined the effects of genetic deletion of the MCU pore-forming subunit of the mitochondrial Ca^2+^ uniporter on tumorigenesis *in vivo* and *in vitro* models. We identified a transformation-associated increase of MCU expression that was associated with enhanced mitochondrial Ca^2+^ uptake and cell proliferation. Genetic deletion of MCU dramatically reduced tumor burden *in vivo*, due to a strong reduction in the proliferative capacity of cells *in vivo* and *in vitro*, particularly under conditions of nutrient stress. Although loss of MCU was associated with enhanced phosphorylation of PDH, mitochondrial respiration was unaffected and pyruvate was diverted to lactate and PDH-independent anaplerosis. In addition, anaplerotic glutamine metabolism was enhanced and agonist-induced Ca^2+^ signaling was altered. Reduced proliferation, delayed cell-cycle progression, increased glycolytic and glutamine metabolism, and a dependence on glucose and glutamine during rapid proliferation suggest that genetic deletion of MCU creates an underlying metabolic defect that strongly inhibits tumorigenesis.

We observed a significant increase in MCU expression as a consequence of cell transformation. To our knowledge, this is the first demonstration of MCU upregulation as a direct consequence of malignant transformation and acquisition of an oncogenic potential. Increased MCU expression resulted in enhanced mitochondrial Ca^2+^ uptake *in vitr*o, in agreement with previous studies (Baughman et al., 2011; De Stefani et al., 2011; Chaudhuri et al., 2013). Hepatic (Ren et al., 2017), breast (Tosatto et al., 2016), and colorectal (Liu et al., 2020) cancers display high expression levels of MCU that positively correlate with tumor size, metastasis, and poor survival prognosis of patients (Curry et al., 2013; Hall et al., 2014). Of note, a worse prognosis in acute myeloid leukemia (Shi et al., 2015), increased proliferation of hepatocellular carcinomas (Guerra et al., 2019), and stronger migratory capacity of breast cancers (Mound et al., 2017) are associated with overexpression of InsP_3_Rs. These observations suggest that enhanced ER-to-mitochondria Ca^2+^ transfer may be a feature associated with many types of cancer. Transformation also facilitated a rapid stimulation of respiration in response to an acute cytoplasmic Ca^2+^signal, which may be crucial for energy regulation to sustain increased bioenergetic demand during cancer progression (Cardenas et al., 2010; Koval et al., 2019; Bustos et al., 2021; Herst et al., 2022).

To determine the role of MCU in tumor progression, we performed subcutaneous xenograft experiments with immortalized and transformed fibroblasts and isogenic HEK293T cells. Cell death was only minimally enhanced by MCU deletion *in vivo*. The absence of significant cell death contrasts with results from our previous *in vitro* studies (Cardenas et al., 2010; Cardenas et al., 2016). A distinction between that study and the present one is the former examined the acute effects of MCU knockdown whereas the present studies used cells with MCU stably deleted. Similarly, acute knockdown of InsP_3_R caused cancer cell death (Cardenas et al., 2010; Cardenas et al., 2016), whereas stable deletion of all InsP_3_R was associated with a significantly repressed proliferative capacity (Young et al., 2022). Compensatory mechanisms have been proposed to account for the lack of overt physiological consequences of MCU-KO in mice of mixed genetic backgrounds (Garbincius et al., 2020). It is likely that metabolic rewiring, such as enhanced nutrient metabolism and switch in TCA substrate preference from glucose to glutamine, as shown here, is a compensatory mechanisms that protect against cell death in stable MCU-KO cell lines used in both our *in vivo* and *in vitro* studies.

In agreement with *Tosatto et. al.* (Tosatto et al., 2016), MCU deletion did not prevent tumor formation but it strongly inhibited tumor growth. Genetic deletion of MCU delayed tumor growth primarily by decreasing cell proliferation that resulted in a smaller tumor size, lower ki-67 index and reduced number of mitotic cells. In addition, lack of MCU was associated with the appearance of giant multinucleated cells, which may reflect quiescent cells that contribute to cancer dormancy. Senescence has been previously associated with reduced transfer of Ca^2+^ from the ER-to-mitochondria (Huang et al., 2000). Furthermore, maintenance of quiescence and escape of hematopoietic stem cells (HSC) from quiescence requires mitochondrial Ca^2+^ uptake (Resende et al., 2010; Umemoto et al., 2018; Ahumada-Castro et al., 2021; Paliwal et al., 2021). Inhibition of cell proliferation by MCU-KO is consistent with anti-proliferative effects of MCU suppression in cancer cells (Cardenas et al., 2016; Ren et al., 2017; Li et al., 2020b; Liu et al., 2020; Wang et al., 2020; Miao et al., 2021; Wu et al., 2021; Zhao et al., 2021), although this has not been universally observed (Curry et al., 2013; Hall et al., 2014; Tosatto et al., 2016; Young et al., 2022). Despite slower tumor growth in xenografts, proliferation *in vitro* of several breast cancer cell lines were independent of MCU (Tosatto et al., 2016), and it has been reported that stable deletion of MCU in HEK293T and HeLa cell lines resulted in enhanced cell proliferation (Young et al., 2022). It is likely that the cell-physiological implications of MCU deletion are cell-type and context-dependent with phenotypes influenced by endogenous metabolic programs (Jose et al., 2011; Pan et al., 2013; Harrington and Murphy, 2015; Luongo et al., 2015; Gui et al., 2016; Marchi et al., 2019; Sullivan and Vander Heiden, 2019). Importantly, the highly-similar effects of MCU deletion in isogenic fibroblasts on cell proliferation *in vivo* and *in vitro* in the present study strongly suggests that mitochondrial Ca^2+^ uptake is essential for cancer cell proliferation, particularly in tumorigenesis.

We found that inhibitory phosphorylation of PDH, the pyruvate gateway to the TCA cycle, is increased in transformed fibroblasts by genetic deletion of MCU and strongly suppressed by MCU rescue. Increased PDH phosphorylation has been consistently observed in response to MCU deletion (Cardenas et al., 2010; Pan et al., 2013; Luongo et al., 2015; Young et al., 2022). The changes we observed are as expected if constitutive mitochondrial Ca^2+^ influx through MCU drives pyruvate dehydrogenase phosphatase (PDP) activity. It is interesting to note that stimulation of Ca^2+^-sensitive dehydrogenases is observed at >500 nM [Ca^2+^]_mit_ (Denton and McCormack, 1980), whereas we observed that basal [Ca^2+^]_mit_ is ∼100 nM in either the presence or absence of MCU. Correlation of the phospho-status of PDH with MCU expression may suggest that PDP is exposed to local high [Ca^2+^]_mit_ that was not resolved in our study. Despite alterations of phospho-PDH, neither basal or maximal respiration were affected by genetic deletion or rescue of MCU in our transformed fibroblasts. Discrepancy between the phospho-status of PDH and mitochondrial respiration suggests that phosphorylation status of PDH in MCU-KO cells may not always faithfully reflect PDH activity. In agreement, our isotope tracing results suggest that flux of carbons from glucose through PDH is not substantially affected by MCU deletion. Glucose-derived m+2 labelled glutamate, fumarate, malate, and aspartate were not different between transformed wild-type and MCU-KO cells, although rescue of MCU enhanced their labeling beyond those observed in the wild-type transformed cells.

The latter observation may suggest that our MCU-KO cells have employed an adaptive metabolic program that promotes TCA-cycle activity independent of MCU-mediated mitochondrial Ca^2+^ uptake. Such an adaptation likely contributes to the observed lack of difference in the OCR between transformed cells with or without MCU. In agreement with our results, strongly enhanced PDH phosphorylation in stable MCU-KO HEK and HeLa cells was not associated with altered glucose-derived carbon flux through PDH or TCA-cycle Ca^2+^-sensitive IDH3 and α-KGDH (Young et al., 2022). PDH is regulated by other factors including elevated NAD^+^/NADH levels observed in MCU-KO cells (Bowker-Kinley et al., 1998; Roche et al., 2003; Patel and Korotchkina, 2006). Elevated pyruvate in chronic MCU-deleted cells could promote sufficient flux through PDH to fuel the TCA cycle at normal rates despite increased inhibitory phosphorylation (St Amand et al., 2000; Spriet and Heigenhauser, 2002). Our observations of enhanced glucose uptake, extracellular acidification and funneling of glucose carbons into lactate suggests that aerobic glycolysis is enhanced in cells lacking MCU. In addition, our results suggest that glucose-derived carbons are funneled into the TCA cycle through alternative pathways involving pyruvate carboxylation in MCU-KO cells. This pathway likely involves PC that generates m+3-labeled fumarate, malate, and aspartate from glucose.

Inhibition of transformed fibroblast proliferation *in vitro* by genetic deletion of MCU was associated with accumulation of the cells in S phase of the cell cycle. It was previously observed that MCU-mediated Ca^2+^ uptake plays a role in mitotic progression (Cardenas et al., 2016) and cell-cycle progression from G1-S phase (Koval et al., 2019), both as a consequence of altered mitochondrial bioenergetics. Our results here suggest that stable deletion of MCU affects mitochondrial metabolism and the production of TCA cycle intermediates that fuel anabolic reactions critical for cell growth and proliferation (DeBerardinis et al., 2007; Mullen et al., 2011; Fendt et al., 2013; Birsoy et al., 2015; Lunt et al., 2015; Cardenas et al., 2016). Generation of biosynthetic substrates by mitochondria supports proliferation by supplementing aspartate for protein and nucleotide synthesis (Birsoy et al., 2015; Sullivan et al., 2015). The accumulation in S-phase observed here is reminiscent of the proliferative phenotype caused by limited aspartate availability for *de novo* synthesis of pyrimidines (Gaglio et al., 2009; Birsoy et al., 2015; Lunt et al., 2015; Saqcena et al., 2015; Sullivan et al., 2015; Patel et al., 2016). Of note, we previously observed that an energetic crisis triggered by acute inhibition or silencing of InsP_3_R or MCU could be rescued by supplementation of exogenous nucleotides (Cardenas et al., 2010; Cardenas et al., 2016; Cardenas et al., 2020). Glutamine provides a nitrogen source for nucleotide synthesis, particularly important to prevent cell-cycle arrest in S phase (St Amand et al., 2000; Patel and Korotchkina, 2006; Saqcena et al., 2015). Thus, glutamine anaplerosis is often a limiting factor for cancer cell growth. Here, we found that deletion of MCU-mediated Ca^2+^ uptake was associated with enhanced glutamine uptake and incorporation of its carbons into TCA-cycle intermediates. We observed a surprisingly high contribution of glutamine for the generation of aspartate pools. In addition, we observed a strong dependence of MCU-KO cell proliferation on glutamine availability. Enhanced lactate production, by fueling NAD^+^ production, may also contribute to enhanced aspartate synstesis by activating the cytosolic malate dehydrogenase to generate oxaloacetate that then drives aspartate synthesis by the aspartate aminotransferase GOT1 (Birsoy et al., 2015).

Despite altered metabolism observed as a consequence of MCU deletion, various bioenergetic parameters, including basal dehydrogenase activity, ROS production, Δψm and [Ca^2+^]_mit_, were independent of MCU in our studies. Lack of effect of MCU deletion on basal bioenergetics has been observed in some cell types (Pan et al., 2013; Luongo et al., 2015; Kwong et al., 2018), although suppression of mitochondrial Ca^2+^ uptake has been reported to disrupt ATP and ROS production and downregulate NAD^+^/NADH ratios in others (Tosatto et al., 2016; Ren et al., 2017; Young et al., 2022). Whereas respiration was unaffected, MCU deletion in our transformed fibroblasts was associated with elevated glycolysis and glutaminolysis, as previously observed in some mouse (Pan et al., 2013; Nichols et al., 2017; Gherardi et al., 2019) and cell (Young et al., 2022) models. An inverse correlation between mitochondrial Ca^2+^ uptake and glycolysis has been observed in epithelial carcinomas and ovarian cancer in which increased MICU1 expression promotes lactate accumulation (Chakraborty et al., 2017; Nemani et al., 2020). Several factors can promote a shift of cellular metabolism towards aerobic glycolysis (Gaude et al., 2018; Szibor et al., 2020; Luengo et al., 2021), including elevated AMPK activity (Young et al., 2022) that we and others previously found were associated with interruption of ER-to-mitochondria Ca^2+^ transfer (Cardenas et al., 2010; Luongo et al., 2015; Ren et al., 2017; Tomar et al., 2019; Zhao et al., 2019; Cardenas et al., 2020). Of note, forced reliance of HEK MCU-KO on the TCA cycle caused a reduction of ATP levels, activation of AMPK and a bioenergetic crisis that led to cell death (Young et al., 2022), similar to the responses of cancer cells to acute deletion of either MCU or InsP_3_R (Cardenas et al., 2016). Furthermore, chronic deletion of MCU in HEK cells results in cell death under conditions in which glycolysis is inhibited (Young et al., 2022). Enhanced glycolysis is broadly associated with proliferation (Diaz-Ruiz et al., 2011), suggesting that enhanced glycolysis in MCU-KO cells may be a compensatory mechanism to sustain cell growth. In previous studies, including our own, acute deletion or knockdown of MCU had significant effects on mitochondrial energy metabolism (Cardenas et al., 2016; Tosatto et al., 2016; Ren et al., 2017; Stejerean-Todoran et al., 2022). However, chronic deletion of MCU has been observed to not result in overt changes of basal mitochondrial metabolism (Pan et al., 2013; Kwong et al., 2015; Luongo et al., 2015; Hamilton et al., 2018; Koval et al., 2019; Wu et al., 2021). Discrepancies between chronic and acute MCU knockout/knockdown models is a recurring theme that might be explained by the activation of compensatory adaptations to maintain mitochondrial energy metabolism (Garbincius and Elrod, 2022). Whereas the apparent flux of glucose into the TCA cycle through PDH was largely unaltered by deletion of MCU, rescue of MCU enhanced this pathway, suggesting that re-wired metabolism in MCU-deleted cells enabled proliferation and survival by engagement of compensatory metabolic pathways.

In addition to identifying a critical role of MCU in tumor growth, we determined that MCU deletion in transformed fibroblasts diminished malignant capabilities *in vitro.* Our studies indicate that inhibition of MCU-mediated Ca^2+^ uptake limits the ability of transformed fibroblasts to form clonally-derived spheres and invade, capabilities associated with initiation and progression of metastatic tumors (Uchida et al., 2010; Ishiguro et al., 2017), as previously observed (Tosatto et al., 2016). Previous studies have suggested a role for MCU as a promoter of invasion and metastasis in breast cancer and colorectal carcinoma-derived cell lines (Curry et al., 2013; Marchi et al., 2013; Tang et al., 2015; Cardenas et al., 2016; Tosatto et al., 2016; Ren et al., 2017; Yu et al., 2017; Liu et al., 2020). Thus, our results suggest that MCU supports cancer malignancy by promoting tumor growth as well as cell-biological functions involved in invasion and recurrence. [Ca^2+^]_cyt_ signaling is a regulator of cell cycle progression (Humeau et al., 2018; Zhao and Pan, 2021). We observed a suppression of agonist-induced InsP_3_R-mediated [Ca^2+^]_cyt_ signals and sustained rise of [Ca^2+^]_cyt_ in MCU-KO fibroblasts. Our findings recapitulate those made by *Koval et.al.*, where genetic deletion of MCU in primary mouse fibroblasts decreased cell proliferation and delayed cell cycle progression through disruption of [Ca^2+^]_cyt_ transients (Koval et al., 2019). MCU helps sustain store-operated Ca^2+^ entry (SOCE) and [Ca^2+^]_cyt_ oscillations by rapid Ca^2+^ buffering, that are expected to be altered by absence of MCU (Kim and Usachev, 2009; Samanta et al., 2014; Zhao and Pan, 2021). [Ca^2+^]_cyt_ dynamics regulate essential processes that promote carcinogenesis and support malignancy. For example, dysregulated cytosolic Ca^2+^ signaling promotes the activation of Ca^2+^sensitive proteins involved in upregulation of epithelial-to-mesenchymal transition, such as calmodulin (Ito et al., 1999; Norgard et al., 2021). Other studies implicate MCU-dependent clearance of cytoplasmic Ca^2+^ and Ca^2+^-sensitive CamKII activation in the regulation of cancer cell proliferation (Koval et al., 2019; Zhao and Pan, 2021), as well as inactivation and nuclear-translocation of the transcription factor NFAT (Kim and Usachev, 2009) implicated in cell survival, angiogenesis, and invasion (Qin et al., 2014). Accordingly, in addition to effects on metabolism, MCU deletion can also perturb cell-cycle progression through the dysregulation of [Ca^2+^]_cyt_ signaling. [Ca^2+^]_cyt_ regulates the activity of the aspartate/glutamate exchangers aralar and citrin, components of the malate-aspartate shuttle (MAS) (del Arco and Satrustegui, 1998; Del Arco et al., 2000; Palmieri et al., 2001; Contreras et al., 2007; Borst, 2020). Of particular importance, activation of the Aralar/MAS pathway promotes regeneration of cytosolic aspartate pools and its inhibition delays cell cycle progression (Contreras et al., 2007; Wang et al., 2016; Alkan and Bogner-Strauss, 2019; Infantino et al., 2019; Del Arco et al., 2023). As noted, cancer cells rely on mitochondrial aspartate for *de novo* pyrimidine synthesis to support uncontrolled proliferation (Brown et al., 2017; Wang et al., 2021), and our isotope labeling experiments revealed enhanced aspartate production from both glucose and glutamine. It is interesting to speculate that slow progression through S phase in MCU-KO cells might result from decreased MAS activity due to MCU-mediated alterations of [Ca^2+^]_cyt_ (Pardo et al., 2006; Patel et al., 2016; Wang et al., 2016; Diehl et al., 2022; Perez-Liebana et al., 2022; Del Arco et al., 2023).

In summary, we discovered that fibroblast cell transformation is associated with upregulation of MCU expression that results in enhanced Ca^2+^ uptake by mitochondria and suppression of inactivating-phosphorylation of PDH that was nevertheless not associated with enhanced pyruvate mtetabolism through PDH or OCR, and enhanced aerobic glycolysis and anaplerotic glutamine and glucose metabolism. Deletion of MCU strongly reduced the tumor burden *in vivo* and decreased cell proliferation *in vitro* and *in vivo*. Mechanistically, this occurred by prolongation of the cell cycle S-phase, a strongly reduced capacity for sphere-forming ability and altered cytoplasmic Ca^2+^ signaling. Our results suggest that targeting MCU may have therapeutic implications in cancer.

## 6 Author contributions

Conceptualization, E.F.G., A.R., K.E.W., Z.A. and J.K.F.; Investigation, E.F.G., M.C.N., C.E.B., J.R.P., and J.S.W.; Writing, E.F.G. and J.K.F. The authors declare no competing financial interests.

## Acknowledgments

We thank Dr. V. Mootha for providing the HEK293T MCU-KO cell line; Dr. R. Payne for discussions and guidance; Dr. T. W. Ridky for advice and lentiviral vectors; Dr. J. Seykora for guidance for histological analyses; the Penn Genomic Analysis Core of the Perelman School of Medicine at the University of Pennsylvania for transformation, template preparation, and purification of plasmids; the Cutaneous Phenomics and Transcriptomics Core (CPAT) for immune staining; the Molecular Pathology and Imaging Core for paraffin embedding, sectioning, and H&E staining of tumor tissue; the Flow Cytometry and Cell Sorting Facility for training and technical assistance.

## 8 Supplementary information

Supplemental information includes 5 figures.

