## Supplementary materials for "The mitochondrial Ca^2+^ channel MCU is critical for tumor growth by supporting cell cycle progression and proliferation"

### *Supplementary Material*

#### 1 Supplementary Figures

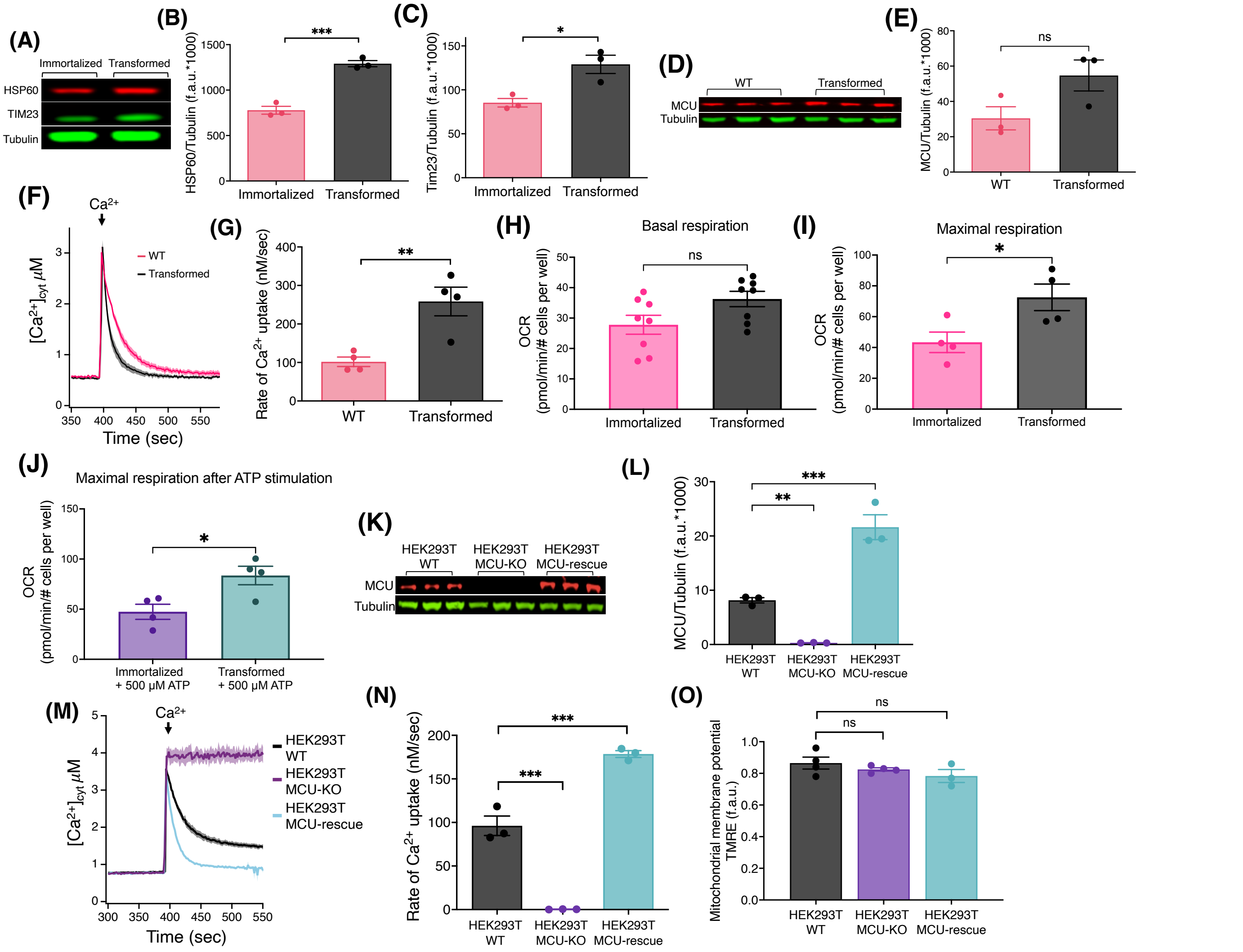


**Supplementary Figure 1.** Transformation of fibroblasts increases mitochondrial mass, upregulates MCU expression and enhances mitochondrial Ca$^{2+}$ uptake. (**A**) Representative immunoblots of HSP60, Tim23 and tubulin in immortalized and transformed fibroblasts. (**B**) HSP60 levels normalized to tubulin detected on the same blots. Values expressed as fluorescence units (f.a.u*1000) (mean ± SEM, n = 3, ***$p < 0.001$, Student's t-test). (**C**) Tim23 levels normalized to tubulin detected on the same blots. Values expressed as fluorescence units (f.a.u*1000) (mean ± SEM, n = 3, *$p < 0.05$, Student's t-test). (**D**) Representative immunoblots of MCU and tubulin in WT and transformed fibroblasts. (**E**) MCU levels normalized to tubulin on same blots. Values expressed as fluorescence units (f.a.u*1000) (mean ± SEM, n = 3, ns = non-significant, Student's t-test). (**F**) Average traces of [Ca$^{2+}$]$_{cyt}$ in suspensions of WT and transformed fibroblasts (mean ± SEM). (**G**) Mitochondrial Ca$^{2+}$ uptake rates of WT and transformed fibroblasts (mean ± SEM, n = 4, **$p < 0.01$, Student's t-test).

(**H**) Basal oxygen consumption rates (OCR) of transformed and immortalized fibroblasts (mean ± SEM, n = 8, ns = non-significant, Student's t-test). (**I**) Maximal uncoupled OCR of transformed and immortalized fibroblasts (mean ± SEM, n = 4, ns = non-significant, Student's t-test). (**J**) Maximal uncoupled OCR of transformed and immortalized fibroblasts after acute stimulation of OCR by ATP (mean ± SEM, n = 4, ns = non-significant, Student's t-test). (**K**) Representative immunoblots of MCU and tubulin in HEK293T WT, MCU-KO, and MCU-rescue cells. (**L**) MCU levels normalized to tubulin expression detected on same blots (mean ± SEM, n = 3, ***p < 0.001, one-way ANOVA). (**M**) Average traces (n = 3) of $[Ca^{2+}]_{cyt}$ in suspensions of permeabilized HEK293T WT, HEK293T MCU-KO, and HEK293T MCU-rescue cells (mean ± SEM). (**N**) Mitochondrial $Ca^{2+}$ uptake rates of HEK293T WT, MCU-KO, and MCU-rescue cells (mean ± SEM, n = 3, **p < 0.01, ***p < 0.001, one-way ANOVA). (**O**) Normalized $\Delta\Psi_m$ (f.a.u) in permeabilized suspensions of HEK293T WT (n = 4), MCU-KO (n = 4), and MCU-rescue (n = 3) cells (mean ± SEM, ns = non-significant, one-way ANOVA).

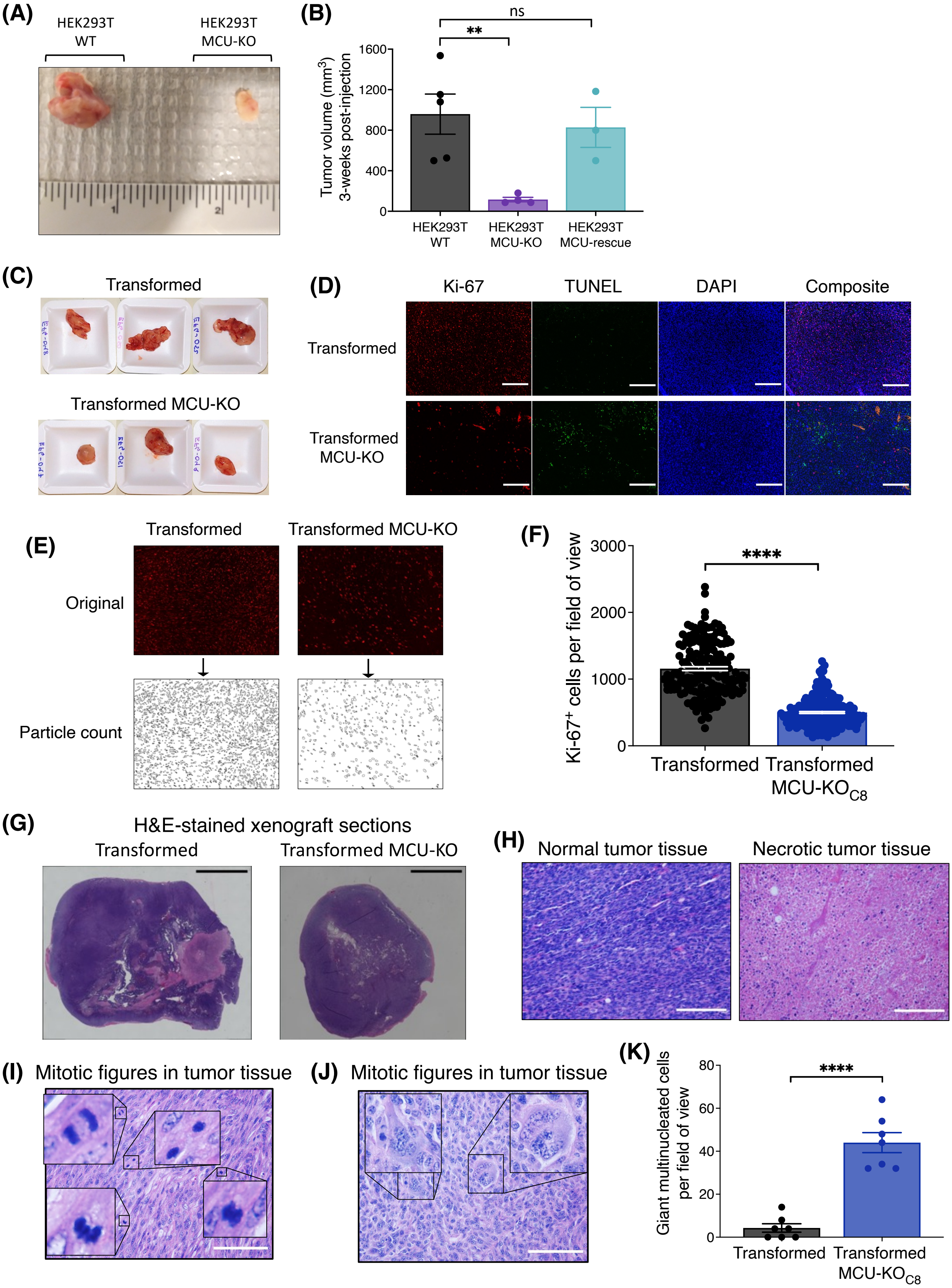
(A)
HEK293T
WT
HEK293T
MCU-KO
(B)
ns
**
Tumor volume (mm3)
3-weeks post-injection
1600
1200
800
400
0
HEK293T
WT
HEK293T
MCU-KO
HEK293T
MCU-rescue
(C)
Transformed
Transformed MCU-KO
(D)
Ki-67
TUNEL
DAPI
Composite
Transformed
Transformed
MCU-KO
(E)
Transformed
Transformed MCU-KO
Original
Particle count
(F)
****
Ki-67+ cells per field of view
3000
2000
1000
0
Transformed
Transformed
MCU-KOC8
(G)
H&E-stained xenograft sections
Transformed
Transformed MCU-KO
(H)
Normal tumor tissue
Necrotic tumor tissue
(I)
Mitotic figures in tumor tissue
(J)
Mitotic figures in tumor tissue
(K)
****
Giant multinucleated cells
per field of view
80
60
40
20
0
Transformed
Transformed
MCU-KOC8

**Supplementary Figure 2.** Image processing and analyses of tumor xenografts. (**A**) Representative tumor xenografts derived from HEK293T WT and MCU-KO cells. (**B**) Volumes of HEK293T WT (n = 5), MCU-KO (n = 4), and MCU-rescue (n = 3) tumor xenografts 3 weeks post-injection (mean ± SEM, $^{**}p < 0.005$, ns = non-significant, one-way ANOVA). (**C**) Representative images of whole tumor xenografts of transformed and MCU-KO fibroblasts. (**D**) Representative immunofluorescence (IF) images of tumor xenograft sections stained for ki-67, TUNEL, and DAPI (scale bar, 200 μm). (**E**) Representative image of ki-67-stained tumor xenograft at magnification of 20x. (**F**) Quantification of ki-$67^{+}$ cells in tumor xenografts of transformed and MCU-KO fibroblasts (mean ± SEM, n = 160, $^{****}p < 0.0001$, Student's t-test). (**G**) Representative images of whole tumor xenografts of transformed and MCU-KO fibroblasts stained with H&E. The whole tumor was imaged at 4x, then all images were stitched together to form a single tumor image (scale bar, 1"; magnification 4x). (**H**) Representative images of hematoxylin and eosin (H&E) stained sections from tumor xenografts emphasizing normal (left) and necrotic (right) tissue (scale bar, 200 μm). (**I**) Representative H&E-stained xenograft section (scale bar, 200 μm) showing mitotic figures delineated by black squares. (**J**) Representative H&E staining of xenograft section emphasizing giant multinucleated cells outlined by black squares (scale bar, 200 μm). (**K**) Quantification of giant multinucleated cells in tumor xenografts. For each tumor, 10 different fields of view were examined as described in Figure 2G (mean ± SEM, n = 7, $^{****}p < 0.0001$, Student's t-test).

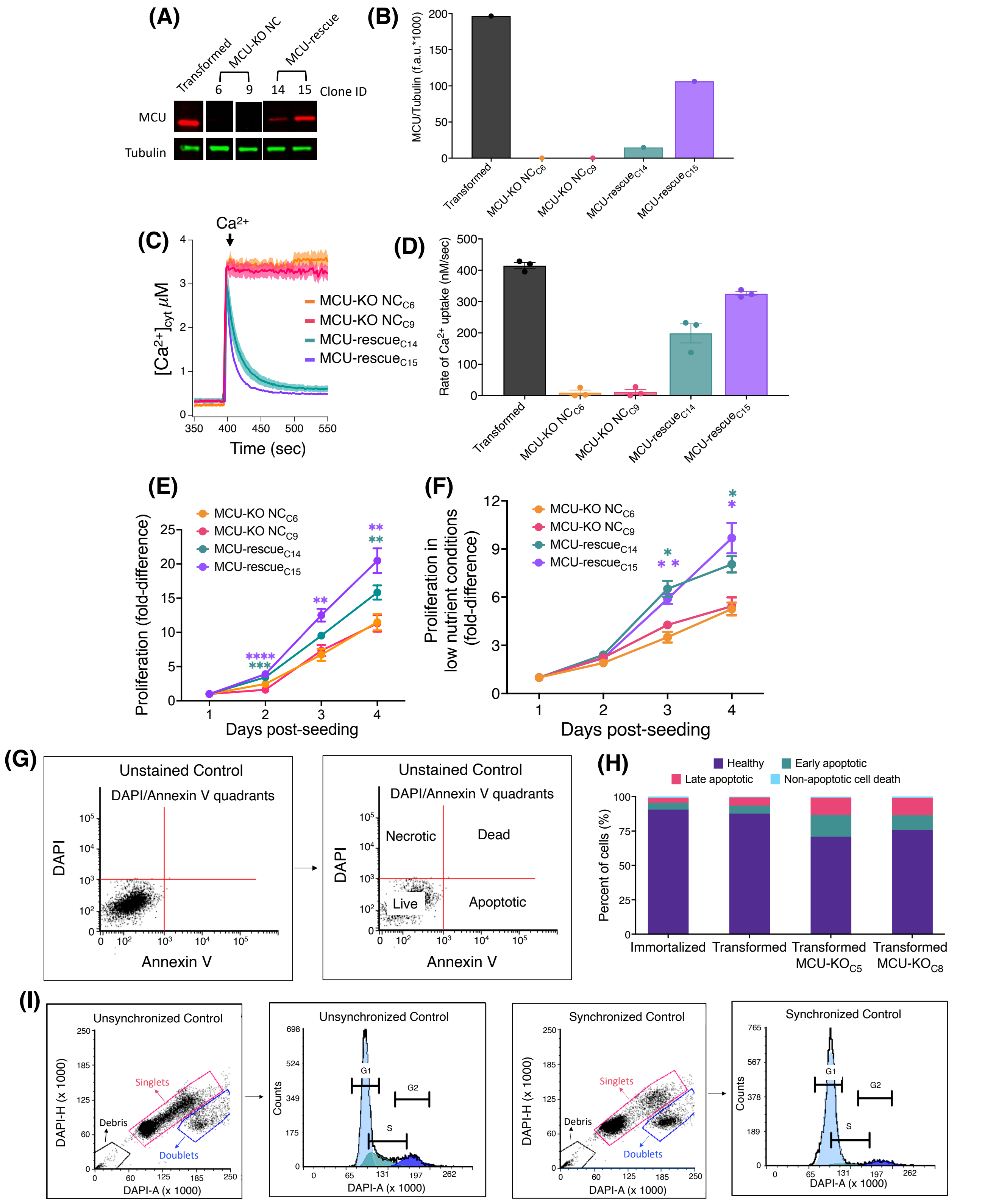


**Supplementary Figure 3.** Rescue of MCU expression increases proliferation of transformed fibroblasts. (**A**) Representative immunoblots of MCU and tubulin in clones of MCU-KO negative controls (NC) and MCU-rescue transformed fibroblasts. (**B**) MCU levels normalized to tubulin. Values expressed as fluorescence units (f.a.u) (n = 1). (**C**) Averaged traces of $[Ca^{2+}]_{cyt}$ in suspensions of MCU-KO NC and MCU-rescue transformed fibroblasts. (**D**) Mitochondrial $Ca^{2+}$ uptake rates of MCU-KO NC and MCU-rescue transformed fibroblasts (mean ± SEM, n = 3, ****p < 0.0001, ns = non-significant, one-way ANOVA). (**E**) Cell proliferation of MCU-KO NC and MCU-rescue clones. Each data point represents 3 biological replicates in triplicate. Fold-difference represents number of cells normalized to day 1 post-seeding. (mean ± SEM, n = 3, **p < 0.01, ***p < 0.001, ****p < 0.0001, two-way ANOVA compared with MCU-KO NC clone 6). (**F**) Cell proliferation of MCU-KO NC and MCU-rescue transformed fibroblasts in low-nutrient conditions. Each data point represents at least 3 biological replicates in triplicate, MCU-KO NC Clone 6 (n = 4), MCU-KO NC Clone 9 (n = 3), MCU-rescue Clone 14 (n = 3), MCU-rescue Clone 15 (n = 4). Fold-difference represents number of cells normalized to day 1 post-seeding. (mean ± SEM, *p < 0.05, **p < 0.01, two-way ANOVA compared with MCU-KO NC clone 6). (**G**) Gating strategy used for the analysis of annexin-V/DAPI flow cytometry data. First, live cells were gated in a forward and side-scatter plot. Then, singlets were gated in a forward-area and forward-height scatter plot. Finally, a 1 µM staurosporine-treated control was used to establish live, dead, and apoptotic quadrants in a DAPI and Annexin V scatter plot. (**H**) Percent (%) healthy cells, early apoptotic cells, late apoptotic/dead cells, and necrotic cells of annexin-V/DAPI FACS plot (mean, n = 3). (**I**) Gating strategy for the analysis of cell cycle flow cytometry data, same gating strategy as in (G). Unsynchronized and 15 µM lovastatin-synchronized samples were used as controls to validate the G1, S, and G2 peaks identified using Multicycle software for cell cycle analysis.

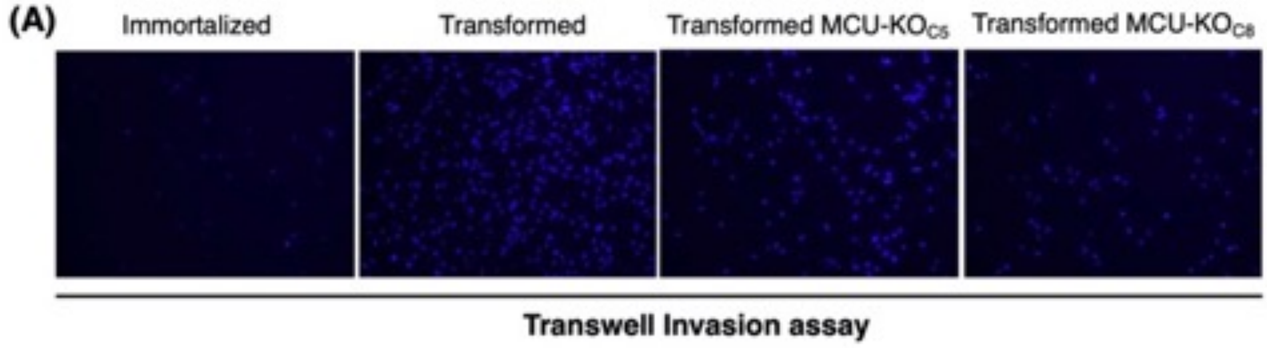


**Supplementary Figure 4.** Matrigel invasion of immortalized and transformed fibroblasts. **(A)** Representative fluorescent images of Hoechst 33342-stained nuclei in Transwell inserts during invasion assay.

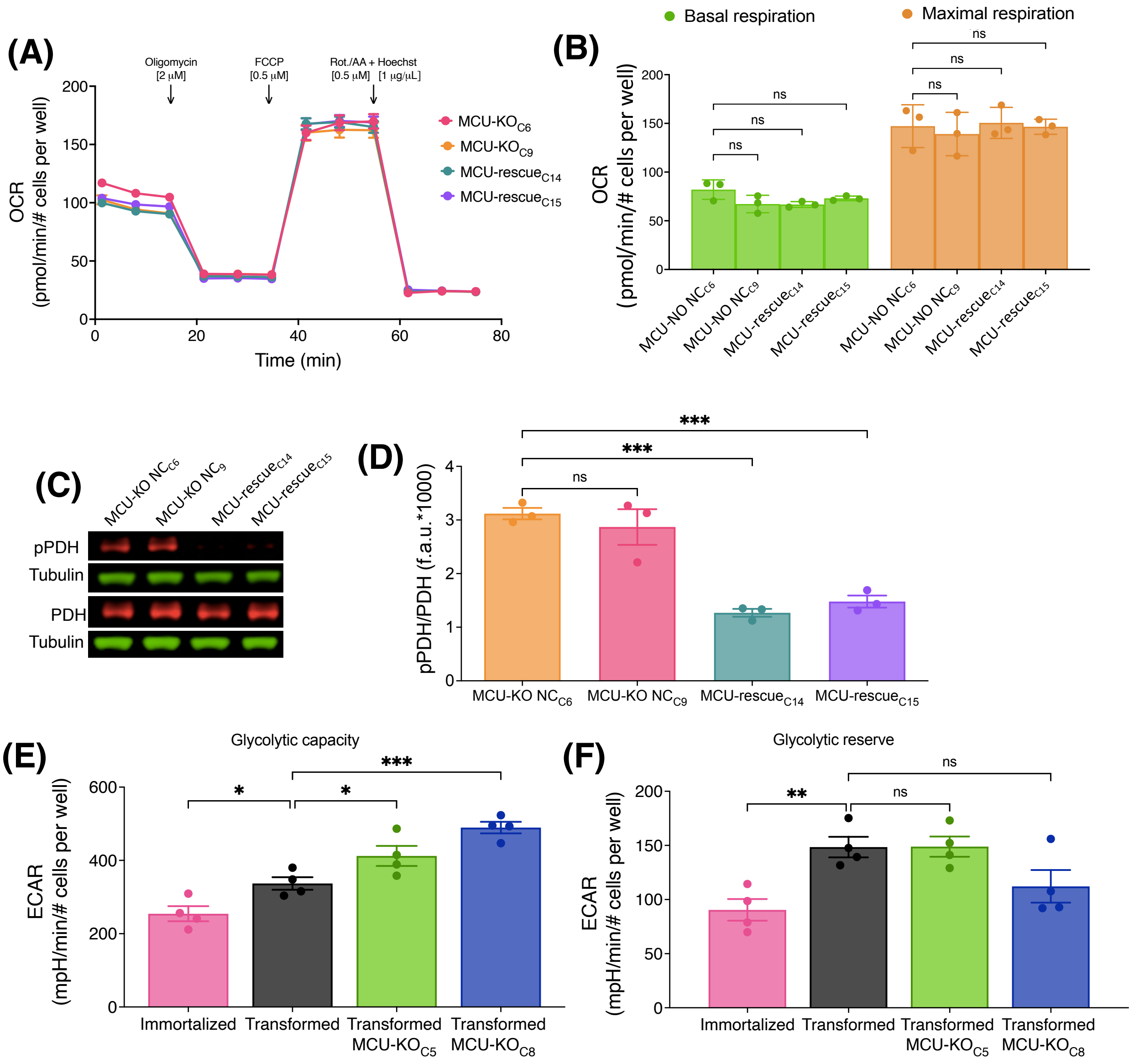


**Supplementary Figure 5.** Effects of MCU deletion on aerobic respiration and glycolysis in transformed fibroblasts. (**A**) Oxygen consumption rates (OCR) of MCU-KO NC and MCU-rescue transformed fibroblasts. Mean values compared to those of transformed fibroblast MCU-KO NC Clone 6 (mean ± SEM, n = 3, two-way). (**B**) Basal (green bars) and maximal (orange bars) respiration of MCU-KO NC and MCU-rescue transformed fibroblasts (mean ± SEM, n = 3, ns = non-significant, two-way ANOVA). (**C**) Representative immunoblots of PDH, pPDH and tubulin in MCU-KO NC and MCU-rescue transformed fibroblasts. (**D**) Relative protein levels of pPDH and PDH determined by measuring intensities of bands normalized to corresponding tubulin band intensity on same blot (mean ± SEM, n = 3, ***$p < 0.001$, ns = non-significant, one-way ANOVA). (**E**) Glycolytic capacity of immortalized and transformed fibroblasts (mean ± SEM, n = 4, *$p < 0.05$, ***$p < 0.001$, one-way ANOVA). (**F**) Glycolytic reserve of immortalized and transformed fibroblasts (mean ± SEM, n = 4, **$p < 0.01$, ns = non-significant, one-way ANOVA).
